# A comprehensive evaluation of generalizability of deep-learning based Hi-C resolution improvement methods

**DOI:** 10.1101/2022.01.27.477975

**Authors:** Ghulam Murtaza, Atishay Jain, Madeline Hughes, Justin Wagner, Ritambhara Singh

## Abstract

**Motivation:** Hi-C is a widely used technique to study the 3D organization of the genome. Due to its high sequencing cost, most of the generated datasets are of coarse resolution, which makes it impractical to study finer chromatin features such as Topologically Associating Domains (TADs) and chromatin loops. Multiple deep-learning-based methods have recently been proposed to increase the resolution of these data sets by imputing Hi-C reads (typically called upscaling). However, the existing works evaluate these methods on either synthetically downsampled or a small subset of experimentally generated sparse Hi-C datasets, making it hard to establish their generalizability in the real-world use case. We present our framework - Hi-CY - that compares existing Hi-C resolution upscaling methods on seven experimentally generated low-resolution Hi-C datasets belonging to various levels of read sparsities originating from three cell lines on a comprehensive set of evaluation metrics. Hi-CY also includes four downstream analysis tasks, such as TAD and chromatin loops recall, to provide a thorough report on the generalizability of these methods.

**Results:** We observe that existing deep-learning methods fail to generalize to experimentally generated sparse Hi-C datasets showing a performance reduction of up to 57 %. As a potential solution, we find that retraining deep-learning based methods with experimentally generated Hi-C datasets improves performance by up to 31%. More importantly, Hi-CY shows that even with retraining, the existing deep-learning based methods struggle to recover biological features such as chromatin loops and TADs when provided with sparse Hi-C datasets. Our study, through Hi-CY framework, highlights the need for rigorous evaluation in future. We identify specific avenues for improvements in the current deep learning-based Hi-C upscaling methods, including but not limited to using experimentally generated datasets for training.

**Availability:** https://github.com/rsinghlab/Hi-CY

**Author Summary:** We evaluate deep learning-based Hi-C upscaling methods with our framework Hi-CY using seven datasets originating from three cell lines evaluated using three correlation metrics, four Hi-C similarity metrics, and four downstream analysis tasks, including TAD and chromatin loop recovery. We identify a distributional shift between Hi-C contact matrices generated from downsampled and experimentally generated sparse Hi-C datasets. We use Hi-CY to establish that the existing methods trained with downsampled Hi-C datasets tend to perform significantly worse on experimentally generated Hi-C datasets. We explore potential strategies to alleviate the drop in performance such as retraining models with experimentally generated datasets. Our results suggest that retraining improves performance up to 31 % on five sparse GM12878 datsets but provides marginal improvement in cross cell-type setting. Moreover, we observe that regardless of the training scheme, all deep-learning based methods struggle to recover biological features such as TADs and chromatin loops when provided with very sparse experimentally generated datasets as inputs.

## 1 Introduction

3D organization of the genome plays a vital role in cell fate and disease onset. A high-throughput chromosome conformation capture experiment, or Hi-C, is a genome-wide sequencing technique that allows researchers to understand and study the 3D organization of the genome [14]. The sequencing results from Hi-C correspond to observed molecular contacts between two genomic loci. This contact information captures local and global interactions of the DNA molecule. In the past decade, analysis of Hi-C data facilitated the discovery of important genomic structural features, including but not limited to A/B compartments [14] that denote active and inactive genomic regions, topologically associated domains (TAD) [3] that represent highly interactive genomic regions, and enhancer-promoter interactions [22] that are involved in the regulation of genes. Therefore, Hi-C experiments are crucial in advancing our understanding of the spatial structure of the genome and its relationship with gene regulation machinery.

When studying the spatial structure of DNA, the quality of the downstream analysis is highly dependent on the resolution of its Hi-C contact map. For example, having precise locations of chromatin loops in a RAD-21 knock-out experiment [21] is crucial in understanding its impact on chromatin structure. Data from a Hi-C experiment coalesces into a matrix (or contact map as shown in Figure 1 **A**) in which rows and columns correspond to fixed-width windows (“bins”) tiled along the genomic axis, and values in the matrix are counts of read pairs that fall into the corresponding bins. The bin size typically ranges from 1 kilobase pair (kbp) to 1 megabase pair (mbp), where the choice of the bin size depends on the number of paired end-reads from the experiment. Lower read count experiments result in sparser contact matrices that require large bin sizes, resulting in a “low-resolution” contact map. Consequently, the downstream analysis of the contact map cannot yield genomic features such as enhancer-promoter interactions that typically occur in the 5 kbp to 10 kbp range [10]. Similarly, the output of a Hi-C experiment with a high number of read counts results in a “high-resolution” contact map with small bin sizes. A contact map of this resolution enables identifying fine-grained genomic features. However, due to the quadratic scaling of the sequencing cost, most tissue and cell line samples do not have any Hi-C data available. For example, on ENCODE portal there are only 92 samples that have a Hi-C experiment available across a collection of 450 cell lines and tissue samples. For cell and tissue samples that have a Hi-C experiment available, they have relatively low read counts (typically *≤* 100 Million reads) and, consequently, low-resolution (*≥* 40 kbp bin size) contact maps [29]. Constructing Hi-C matrices with high resolution (*≤* 10 kbp bin size) requires billions of Hi-C reads [10], which can be prohibitively expensive to obtain for many experiments. Thus, the absence of such matrices makes the comprehensive analysis of the spatial structure of DNA difficult. This limitation is even more apparent in single-cell variants of Hi-C protocol [19], where the reads are even more sparse, and it is a complicated experimental challenge to acquire high-resolution contact matrices.

**Figure 1:**
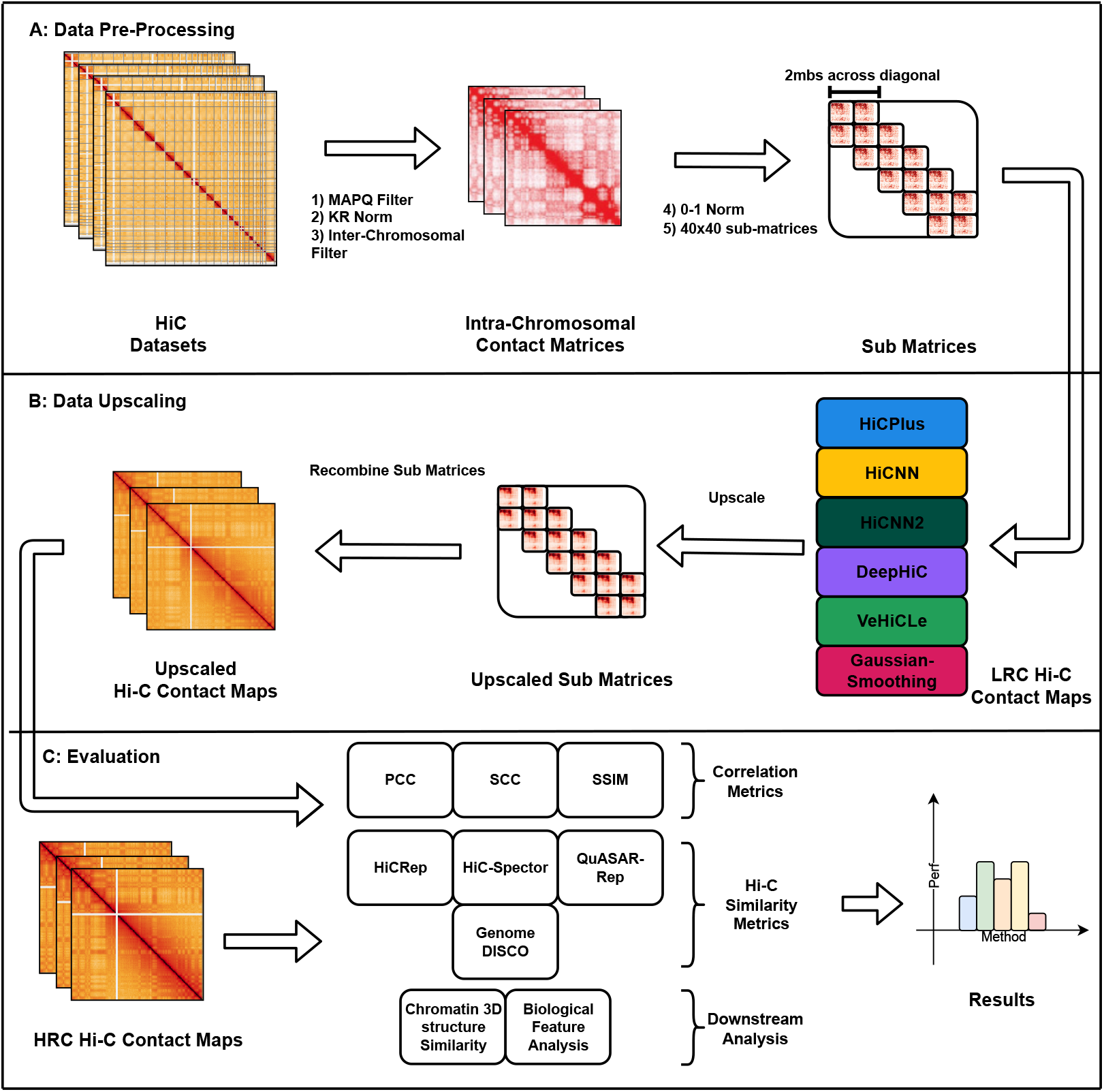
Overview of our benchmarking framework Hi-CY. **A** Data pre-processing pipeline - We: 1) Filtered the Hi-C matrices with the same MAPQ value (*≥*30). 2) Normalized them with the same KR normalization algorithm to ensure a fair comparison. 3) We removed inter-chromosomal contacts because of their extremely sparse nature. 4) Performed a 0-1 normalization on intra-chromosomal matrices to reduce the impact of extreme values. 5) We cropped appropriately sized sub-matrices to ensure that input is in the correct format for each upscaling algorithm. **B** We upscaled the sub-matrices using a wide variety of deep learning-based upscaling models (and Gaussian Smoothing) and then recombined them to form upscaled intra-chromosomal Hi-C matrices. **C** We combined multiple Hi-C similarity metrics, correlation-based metrics and downstream analyses, like chromatin loops and TAD recovery analysis, to provide a comprehensive report that we can use to analyze the performance of each upscaling model.

Recently, researchers have developed several computational methods to upscale^1^ Hi-C matrices by imputing Hi-C reads. Deep learning-based methods [8, 15–17, 29] have shown remarkable success in upscaling Hi-C matrices when trained and tested on simulated datasets. These datasets are typically constructed by uniformly removing a fixed fraction of reads from an experimentally generated high-resolution Hi-C contact map to simulate a low-resolution (downsampled) Hi-C contact map. This downsampling method tends to miss out on experimental artifacts such as diagonal effect [2] and produces Hi-C contact map that have substantially different distribution from a experimentally generated sparse Hi-C contact map. Moreover, the existing methods report their performance on correlation-based metrics that ignore the genome’s multi-scale hierarchical organization. VeHiCle is an exception that [7] utilizes experimentally generated Hi-C datasets and state-of-the-art Hi-C-specific similarity metrics [27]. VeHiCle restricts its analysis to only four datasets with low sparsity (high number of reads) Hi-C datasets. Therefore, it does not provide a comprehensive report on the utility of the method on in its intended use case, where Hi-C matrices tend to be more sparse.

We investigate how existing deep-learning based methods generalize their upscaling performance to experimentally generated sparse Hi-C datasets. Given the rising popularity of single-cell chromatin capture protocols such as scHi-C [19], that are even sparser than their bulk sequencing counterparts, its imperative to test how these methods perform for sparse Hi-C datasets for future methods development. To accomplish this, we developed a framework to pre-process, upscale, and evaluate methods on multiple Hi-C contact maps with varying levels of read counts originating from different cell lines. We present our framework, Hi-CY, in Figure 1. Hi-CY supports the investigation of Hi-C resolution upscaling efforts by providing access to a comprehensive set of Hi-C datasets and evaluation metrics. We contribute towards the recent efforts to standardize evaluation protocols and develop reproducible benchmarks such as those for epigenome imputation [23], protein-drug interaction prediction [9], and Hi-C feature extraction [4]. These projects perform a comprehensive evaluation of existing methods and provide robust and reproducible frameworks to make them accessible and accelerate research.

Hi-CY includes seven sparse experimentally generated datasets from three cell lines (GM12878, K562, and IMR90) with sparsity ranging from a 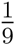 to 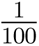 of reads in comparison to the appropriate high-resolution Hi-C contact map. We also package six Hi-C upscaling methods (Gaussian Smoothing, HiCPlus, HiCNN, HiCNN2, DeepHiC, and VeHiCLe) that we evaluate on three correlation-based metrics, four Hi-C similarity metrics, and four downstream analysis tasks. Our metrics capture the bin-wise similarity (through correlation-based metrics), genome structural similarity, and biological information content to provide a comprehensive report on the utility of these generated Hi-C contact maps. Our results show:

- The existing deep-learning based methods struggle to generalize to sparse experimentally generated Hi-C inputs. These methods show a reduction of up to 57% in performance when upscaling a Hi-C contact map with 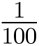 of the reads (compared to the high-resolution Hi-C contact map) in comparison to the upscaling a Hi-C contact map with 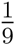 of the reads.
- Our results show that retraining existing deep-learning based models with experimentally generated Hi-C datasets improves performance by upto 31% on experimentally generated Hi-C contact maps.
- We further find that deep-learning based methods still struggle to recover biologically meaningful features in the upscaled Hi-C contact maps, even when retrained, particularly on chromatin loop and DNA hairpin recovery tasks.

## 2 Hi-CY: A comprehensive evaluation framework

We developed Hi-CY to facilitate developing and evaluating Hi-C upscaling methods in a robust and reproducible setup. As shown in Figure 1, Hi-CY packages: 1) a Hi-C pre-processing pipeline, 2) a Hi-C upscaling pipeline with five deep-learning based methods, and 3) a comprehensive evaluation pipeline under a unified framework. Our code is available publicly on GitHub at https://github.com/rsinghlab/Hi-CY.

### 2.1 Hi-C pre-processing

After reviewing existing deep-learning based upscaling methods, [7, 8, 15–17, 29] we use Hi-C experiments from GEO Accession GSE63525 for GM12878, IMR90, and K562 as our primary high resolution that we refer to as High Read Count^2^ (HRC) datasets [10]. Similar to previous evaluation [8, 15, 16, 29], we generate 12 downsampled datasets by uniformly downsampling primary GM12878, IMR90, and K562 datasets by a factor of 16, 25, 50, and 100. We collect an additional seven experimentally generated Low Read Count (LRC)^3^Hi-C datasets to evaluate performance in real-world settings on sparse matrices. Five of these LRC datasets are for the GM12878 cell line and have sparsity ranging from 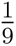 to 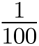 reads compared to the HRC dataset. The remaining two LRC datasets are for IMR90 and K562 cell lines with 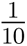 and 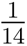 of reads, respectively. We also include a GM12878 HRC biological replicate cell line [10] in our analysis to calculate the “upper-bound” on the metric performance, which we show as a black dotted line wherever appropriate. We show absolute read counts, sparsity, and the source experiment of all datasets in Table 1.

**Table 1:** Summary of the datasets and their sources. All experiments used Mbol enzyme and filtered fragments to be in the size range of 300-500 using SPRI beads. HRC refers to high read count and LRC refers to low-read-count Hi-C contact matrices. Sparsity represents the fraction of reads in comparison to the relevant HRC Hi-C contact map.

| Dataset | Absolute Read Counts | Sparsity | Source |
| --- | --- | --- | --- |
| GM12878-HRC-1 | 1,844,107,778 | 1 | GEO (GSE63525) |
| GM12878-HRC-replicate | 1,564,534,654 | N/A | GEO (GSE63525) |
| GM12878-LRC-1 | 42,453,795 | 1/44 | ENCODE (ENCSR382RFU) |
| GM12878-LRC-2 | 37,079,587 | 1/50 | ENCODE (ENCSR382RFU) |
| GM12878-LRC-3 | 70,138,184 | 1/25 | ENCODE (ENCSR968KAY) |
| GM12878-LRC-4 | 202,380,884 | 1/9 | GEO (GSM1551575) |
| GM12878-LRC-5 | 18,696,952 | 1/100 | GEO (GSM1551582) |
| IMR90-HRC-1 | 735,043,093 | 1 | GEO (GSE63525) |
| IMR90-LRC-1 | 75,193,876 | 1/10 | GEO (GSM1551606) |
| K562-HRC-1 | 641,402,880 | 1 | GEO (GSE63525) |
| K562-LRC-1 | 44,882,605 | 1/14 | GEO (GSM1551622) |

For these Hi-C contact maps, we pre-process them by filtering reads using a MAPQ value of *>*=30 and performed KR normalization to remove reads with low statistical confidence as well as account for experimental artifacts. We sampled both LRC and HRC datasets at 10Kbp resolution and only kept intra-chromosomal contacts to generate twenty-two dense contact matrices. We performed 99.95th percentile normalization to scale all observed contacts between 0 and 1, which has been shown to improve the predictive capabilities of deep-learning based models [8]. Additionally, we cropped out sub-matrices (size depending on each model’s input parameters) across a 2 Mbp range of the diagonal, as this is shown to contain contacts with the highest biological information content [29]. Finally, we divided the data for 22 chromosomes into training (chr1-chr8 and chr12-chr18), validation (chr9-chr12), and test (chr19-chr22) sets. We exclusively used GM12878 datasets to train our models, as is the standard for all methods we compare.

### 2.2 Deep-learning based Hi-C upscaling models

We set up five state-of-the-art deep-learning based Hi-C upscaling methods divided into two broad categories – those that employ an adversarial loss to optimize weights and those that do not. HiCPlus [29] uses a three-layer convolutional network to optimize with Mean Squared Error (MSE) loss. HiCNN [16] extends the HiCPlus approach to a 54-layer architecture that uses residual connections between the CNN layers [6], improving the performance, training times, and stability. HiCNN2 [17] ensembles HiCNN and HiCPlus with a VDSR (Very Deep Super Resolution) model to generate an output that is a linear combination, with learned weighted contribution, of all the networks. HiCGAN [15], DeepHiC [8], and VeHiCLe [7] employ Generative Adversarial Networks. GANs [5] jointly optimize two models – a generator that produces samples and a discriminator that tells fake samples from real ones – to learn to create increasingly more realistic outputs. HiCGAN uses an MSE loss and cross-entropy (computed through discriminator output) to optimize the weights. DeepHiC extends HiCGAN to introduce a perceptual and total variation loss to generate Hi-C contact maps with sharper and more realistic features. Lastly, VeHiCle makes two modifications to the DeepHiC approach: 1) It replaces the perceptual loss with a domain-specific Hi-C loss using an unsupervised model trained to generate Hi-C input, and 2) It adds insulation loss, forcing the model to learn the underlying biological structure (specifically topologically associated domains) and generate more informative Hi-C matrices.

For our evaluations, we retrain HiCPlus, HiCNN, and HiCNN2 to ensure these models generate an output of value between [0, 1]. These three methods predict raw contact counts, making the generated Hi-C matrices incomparable with GAN-style methods and putting these models at a disadvantage against other methods, as shown by DeepHiC [8]. Similar to DeepHiC, we train four different versions, each on 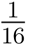, 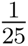, 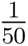 and, 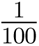 downsampled datasets to provide a range of Hi-C input sparsities [8]. When upscaling, we utilize the version with the smallest difference between the sparsity of input Hi-C data and the downsampled Hi-C data we used to train the method. We show the training curves in the Supplementary Section S1. Note that we exclude HiCGAN [15] from our comparisons as we could not set up the provided code base as it depended on deprecated Python packages. For DeepHiC and VeHiCle, we used the provided weights. As a baseline upscaling algorithm, we added gaussian smoothing with a kernel size of *n* = 17 and 2D Gaussian distribution with *σ_x_* = *σ_y_* = 7, we found these parameters to provide best performance. We keep the architecture the same for all the retrained methods. We do not tune layer parameters to ensure we compare the off-shelf versions of these methods to test their real-world applicability. Lastly, we use the same merging algorithms provided by these methods to combine the predicted samples into full chromosome contact maps.

### 2.3 Evaluation metrics

Recent work has shown that correlation metrics, such as Pearson’s Correlation Coefficient (PCC) and Spearman’s Correlation Coefficient (SCC), fail to assign an accurate reproducibility score for Hi-C experiments due to a limitation in accounting for the underlying data distributions (e.g., distance effect in Hi-C matrices) [27]. Unfortunately, most of the current Hi-C upscaling studies have used PCC and SCC along with Structural Similarity Index Metric (SSIM), which calculates similarity using two images’ luminance, contrast, and structure to evaluate their performance. Due to the limitations of these correlation-based metrics, we include the Hi-C similarity metrics [27] - GenomeDISCO, HiCRep, Hi-C-Spector, and QuASAR-Rep - in our evaluation pipeline. We show results for GenomeDISCO as our main metric, which uses random graph walks for smoothing out the contact matrices at varying scales to compute concordance scores between the two input maps. Similarity across these smoothed contacts corresponds to similarity at various genomic organizational scales [24]. We show results for the rest of the metrics (and explain their mechanisms to compute similarity) in the Supplementary Section S3. For all of the reported metrics, a higher score (closer to 1) is better.

### 2.4 Downstream analyses for biological validation

For a biologically informed evaluation, we perform four additional downstream analyses to assess the utility of information recovered from the upscaled Hi-C matrices. While Hi-C similarity scores produced by - GenomeDISCO, QuASAR-Rep, HiC-Rep, and HiC-Spector - are developed to compare the higher-order structure of the chromatin that can be recovered through Hi-C contact maps, they do not compare biological features such as TADs and chromatin loops. Therefore, we evaluate the quality of 3D reconstruction, recovered chromatin loops, TADs, and DNA hairpains from upscaled Hi-C maps to provide a thorough analysis on the utility of the predicted Hi-C contact maps in their real-world downstream use case.

#### 2.4.1 Quality of chromatin reconstruction from upscaled Hi-C maps

We use 3DMax [20] to generate a 3D model of chromatin from both the HRC and the upscaled Hi-C matrices. 3DMax uses iterative correction with eigen-decomposition to denoise and normalize the Hi-C contact map before constructing the 3D model. We use default 3DMax parameters and compare the 3D models using the Template Modeling score (TM-score), which is typically used in protein-structure comparison, to estimate the reconstruction accuracy. A score closer to 1 indicates a more similar 3D structure, that in our case corresponds to having similar genome organization and function. Typically, a score greater than 0.5 suggests that 3D models have similar underlying structures [11].

#### 2.4.2 Quality of recovered chromatin loops, TADs, and DNA hairpins from the upscaled Hi-C maps

To analyze the structural features of the genome, we used Chromosight [18] on Hi-C contact matrices to call chromatin loops, TADs, and DNA hairpins. Recent works have shown these features are crucial in understanding gene regulation, disease, and genome organization [1, 10, 22]. We recovered these features from the Hi-C matrices using Chromosight [18] with its default parameters and detection kernels. Given Chromosight’s ability to recover an extensive set of biologically informative features [18], we consider the chromatin features retrieved from the HRC matrices as a “ground truth” feature set and compare the derived features from the upscaled matrices against them. We count the following: 1) True Positives (TP) that overlap^4^ in both matrices; 2) False Positives (FP), features that are called on the imputed matrices but are not present in the HRC matrices; and 3) False Negatives (FN), features in HRC matrices that were absent in imputed matrices. Then we compute the F1 score using:

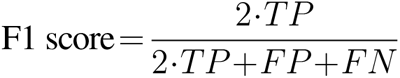

## 3 Results

In this section, we detail the results of the following experimental steps: 1) reproduce previous results using our Hi-CY framework, 2) use Hi-CY to show that deep-learning based methods’ performance does not generalize to experimentally generated LRC datasets, 3) characterize the potential issues leading to poor generalization, and 4) present potential strategies to improve the generalization of these methods on experimentally generated LRC datasets. For all presented results, we show region chr22:41-43Mbp to qualitatively visualize feature recovery performance. We picked this region because it contains a high density of chromatin features including TADs, chromatin loops, and DNA hairpins. The quantitative evaluations show averaged cross-chromosome scores on all four (chr19-chr22) test chromosomes for all metrics and downstream analyses.

### 3.1 Hi-CY framework reproduces similar performance of deep learning based models on downsampled datasets

All existing deep learning-based Hi-C resolution upscaling models [8, 15–17, 29] show that they can achieve correlation and reproducibility scores comparable or better than a biological replicate when upscaling a downsampled Hi-C contact map. These methods show their performance on Hi-C contact maps with a downsampling ratio of up to 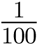 reads compared to the output HRC contact map [8]. We reproduce performance trends using Hi-CY to establish the reproducibility and accuracy of our approach. We generate 12 Hi-C datasets for three cell lines - GM12878, IMR90, and K562 - downsampled by factors of 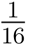, 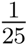, 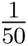 and 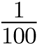 of the reads in the corresponding HRC matrix. We evaluate the performance by comparing the generated Hi-C contact maps against the HRC contact maps. In Figure 2 **A**, we visualize the predicted Hi-C contact maps when using an input GM12878 contact map downsampled by a factor of 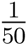 and compare them against the target HRC GM12878 dataset. All methods, excluding the baseline Gaussian-Smoothing, can recover the finer features, including the sub-TAD structures highlighted by a blue dotted rectangle, with DeepHiC generating the most visually similar Hi-C matrices compared to the target HRC contact map.

**Figure 2:**
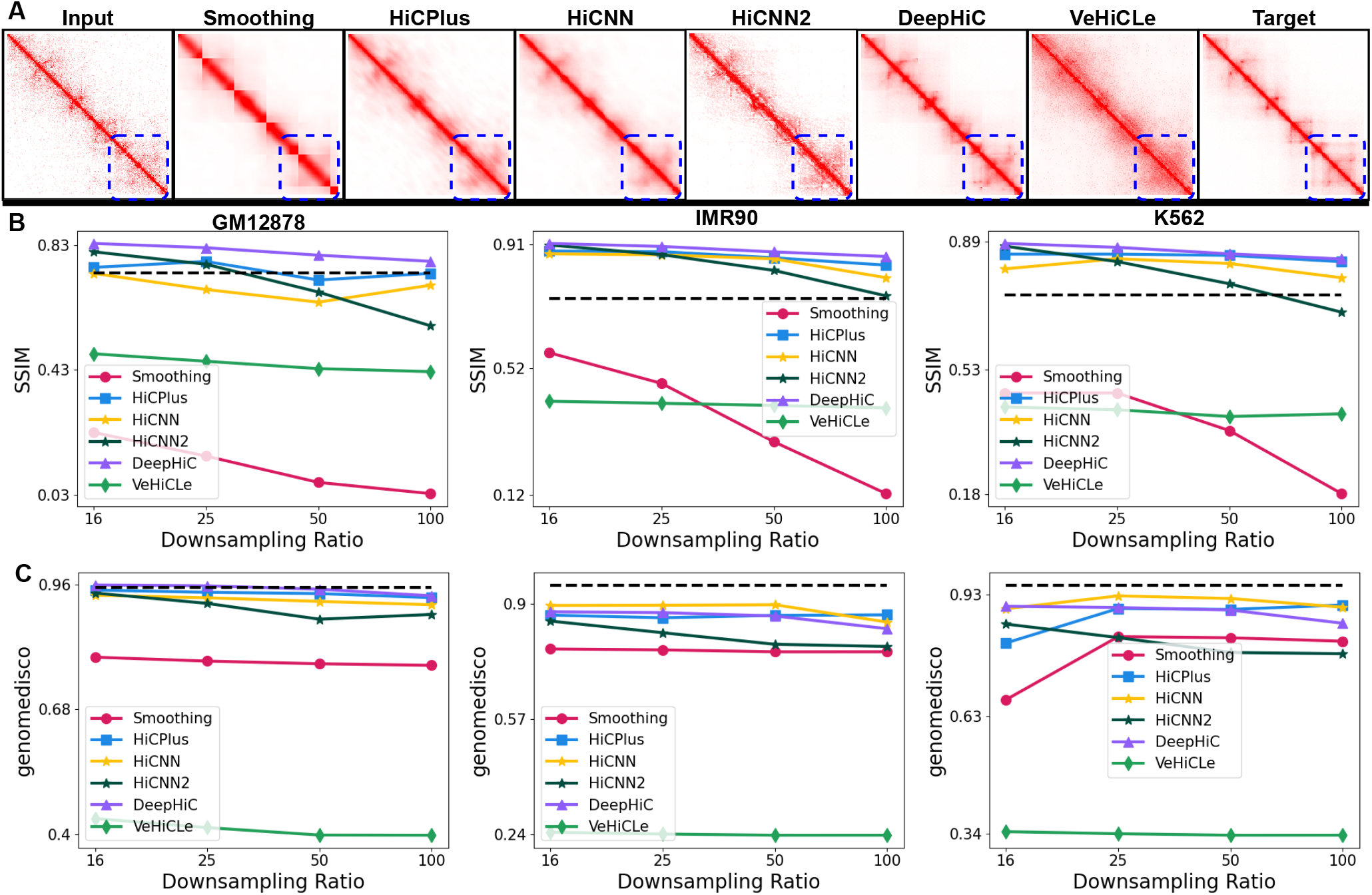
Hi-CY can reproduce the performance of deep learning-based methods on upscaling downsampled input Hi-C datasets. **A** Deep learning-based methods’ output for chr22:41-43Mbp with input reads downsampled to 1/50 counts of the original HRC Hi-C matrix. **B-C** On the x-axis, we present the downsampling ratio in increasing order, and on the y-axis, we report SSIM (panel **B**) and GenomeDISCO (panel **C**). As the downsampling ratio increases, HiCPlus, DeepHiC, HiCNN, and HiCNN2 show similar or comparable performance compared to the biological replicate shown as a dotted black line. VeHiCle and Gaussian Smoothing baseline give a lower performance on these datasets.

We quantify the performance in Figure 2 **B-C** by comparing SSIM and GenomeDISCO scores (on the y-axis) across all three cell lines and four different downsampling ratios (on the x-axis). Our results demonstrate that DeepHiC, HiCNN, HiCNN2, and HiCPlus perform better than or comparable to the biological replicate (shown as a dotted black line) on SSIM and GenomeDISCO metrics. Moreover, these methods generalize to both downsampling ratios and cross-cell type inputs. In contrast, VeHiCLe offers consistent but lower performance than other methods on both metrics. We expected this result given VeHiCle was trained and evaluated on a experimentally generated LRC Hi-C dataset (GSE63525 HIC0001) [7]. These results indicate Hi-CY can train and evaluate the existing methods in a unified pipeline and reproduce results from the previous studies.

### 3.2 Testing generalization of deep-learning based models on experimentally generated Hi-C datasets

We use the Hi-CY framework to evaluate the performance of existing models on experimentally generated LRC Hi-C datasets and contrast to contrast it with their performance on downsampled Hi-C datasets we used in the previous experiment. We test generalizability on five datasets from the GM12878 cell line with read count sparsity ranging from 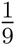 to 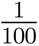 of the total reads of the GM12878 HRC Hi-C matrix. We further test one dataset each from K562 and IMR90 with a sparsity of 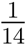 and 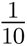 respectively, to evaluate cross-cell-type generalization. First, we compare the output produced by the models when provided with the GM12878-LRC-4 dataset with 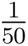 of the reads of the GM12878 HRC Hi-C contact map. As shown in the Figure 3 **A** all methods, including VeHiCLe, struggle to recover chromatin features, particularly the sub-TAD features highlighted with a dotted blue rectangle, which they could recover when provided a downsampled contact map with a similar number of reads. Next, as shown in Figure 3 **B**, we observe that none of the methods can get scores similar to the biological replicate (shown with the dotted black line). Moreover, except VeHiCLe, all methods show a decrease in performance as the sparsity increases. For example, comparing HiCNN results on GM12878 datasets with 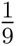 to 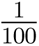 of the reads, SSIM goes from 0.77 to 0.45 while GenomeDISCO goes from 0.94 to 0.4. Although VeHiCLe shows stable scores, they are lower than other methods for SSIM and GenomeDISCO, suggesting it might have overfit to the dataset it was trained with (GSE63525 HIC0001).

**Figure 3:**
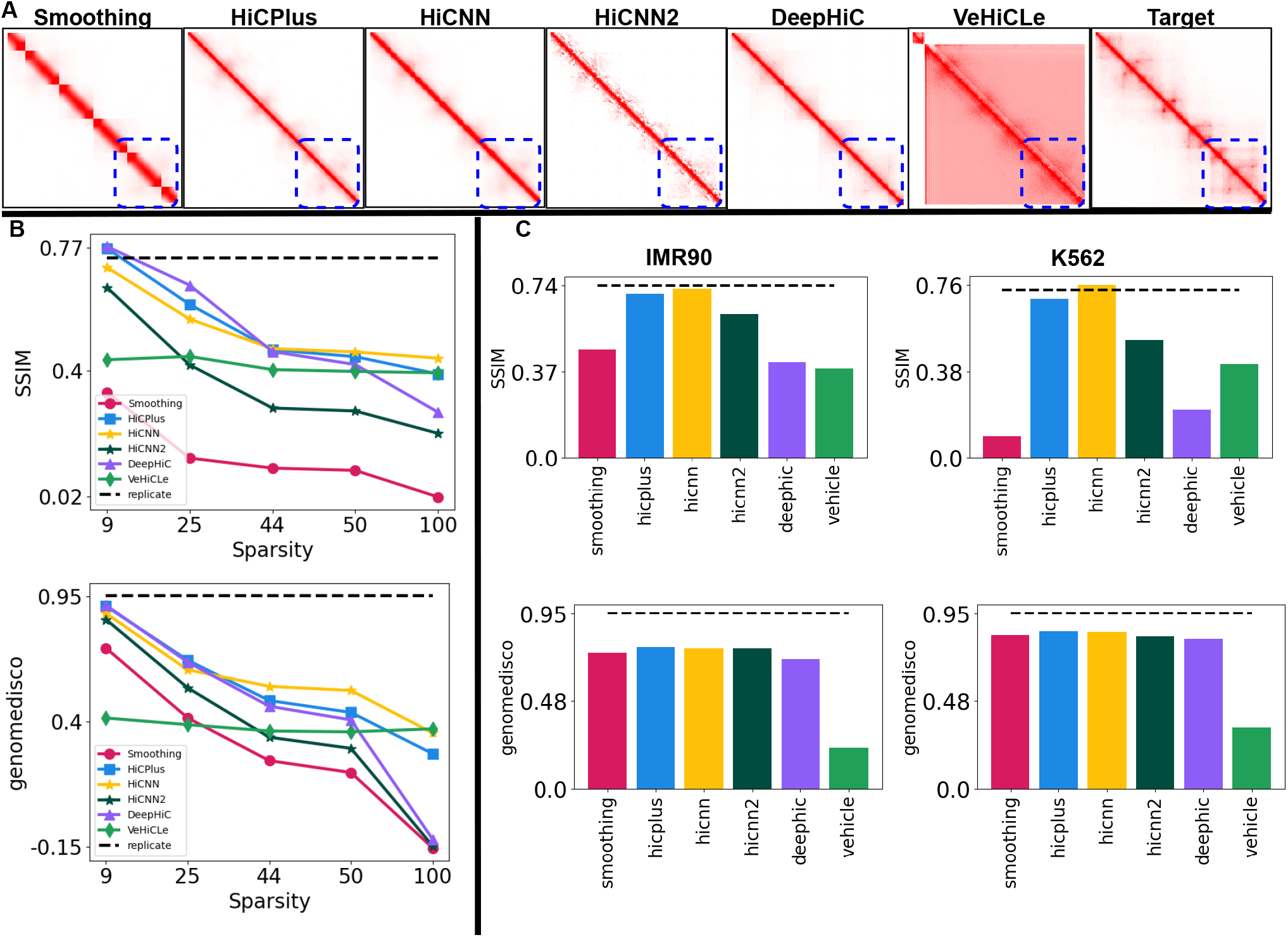
Performance comparison for upscaling experimentally generated LRC Hi-C datasets. **A** We visualize the outputs of our methods on the GM12878-LRC-2 dataset with a read count of 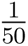 of the total reads of GM12878 HRC Hi-C map for chr22:41-43Mbp shows that all methods struggle to recover finer chromatin features in comparison to when provided with a downsampled Hi-C contact map. **B** We quantify the decrease in performance of upscaling methods as we increase the sparsity (on the x-axis) of GM12878 LRC datasets on SSIM and GenomeDISCO metrics (on the y-axis). We observe that all methods show a substantial drop in performance as we increase sparsity, with HiCNN showing better robustness to the sparsity of data in comparison to other methods. **C** We show a drop in the performance of these methods on IMR90 and K562’s LRC datasets by comparing them against a biological replicate score shown with a dotted black line. We observe a substantial drop in performance in both cell lines on GenomeDISCO metrics. Our results (**B-C**) show that all methods fail to generalize to sparse experimentally generated Hi-C datasets.

Lastly, we test how these deep-learning based models generalize to experimentally generated cross-cell type sparse Hi-C contact maps. Figure 3 **C** shows the SSIM and GenomeDISCO scores on the upscaled matrices when provided with IMR90-LRC-1 and K562-LRC-1 datasets as inputs. Across IMR90 and K562 cell lines, both HiCNN and HiCPlus achieve SSIM scores similar to the biological replicate. However, on the GenomeDISCO metric all methods show a substantial reduction in performance across both cell lines. Our results show that all methods strongly favor downsampled datasets and perform significantly worse on experimentally generated sparse LRC Hi-C datasets. Our results on the rest of the metrics are presented in the Supplementary Figures S7, adding further support our findings. Overall, among the available Hi-C upscaling methods, HiCNN shows the best generalizability to its real-world use cases out-of-the-box with a competitive performance on GM12878 datasets (particularly highly sparse 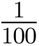 dataset) and the best performance on IMR90 and K562 datasets.

### 3.3 A significant distributional shift between downsampled datasets and experimentally generated LRC datasets hurts the generalizability of deep learning models

Lack of generalizability in deep-learning models is typically attributed to distributional differences [13]. We compare the distribution of reads in LRC and downsampled Hi-C datasets and the impact of that difference on the underlying chromatin structure. First, we compare the observed reads between an experimentally generated LRC Hi-C contact map and a downsampled contact map with similar reads. In Figure 4**A**, we plot the log (base 10) difference of region chr22:41-43Mbp between experimentally generated GM12878 LRC dataset and downsampled datasets with 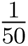 of the reads of the HRC Hi-C contact map as a heat map. The heatmap highlights the region with higher log differences in yellow compared to regions with smaller differences in purple. We observe bins with a higher difference on the heatmap as we move further away from the diagonal. To establish this trend across all the chromosomes (test, train, and validation), we plot PCC between observed reads in experimentally generated or downsampled and HRC datasets at increasing genomic distances. HiCNN, HiCPlus, and DeepHiC [8, 16, 17, 29] have used this analysis to compare the similarity and distribution of Hi-C samples across each genomic distance. In this curve, a higher correlation on all genomic distances suggests a higher read count distribution similarity between the two samples. Figure 4 **B** shows that across all genomic distances, the experimentally generated LRC Hi-C dataset has a lower correlation than the downsampled datasets, with the difference between those two increasing as the distance increases. These observations suggest that experimentally generated LRC Hi-C contact maps have a higher distributional difference with HRC Hi-C contact maps than downsampled Hi-C contact maps with a similar number of reads. This difference arises because uniformly removing reads to downsample Hi-C contact matrices fails to account for the distance-effect [2], suggesting that Hi-C reads are more likely to be observed on genomic bins closer to the diagonal.

**Figure 4:**
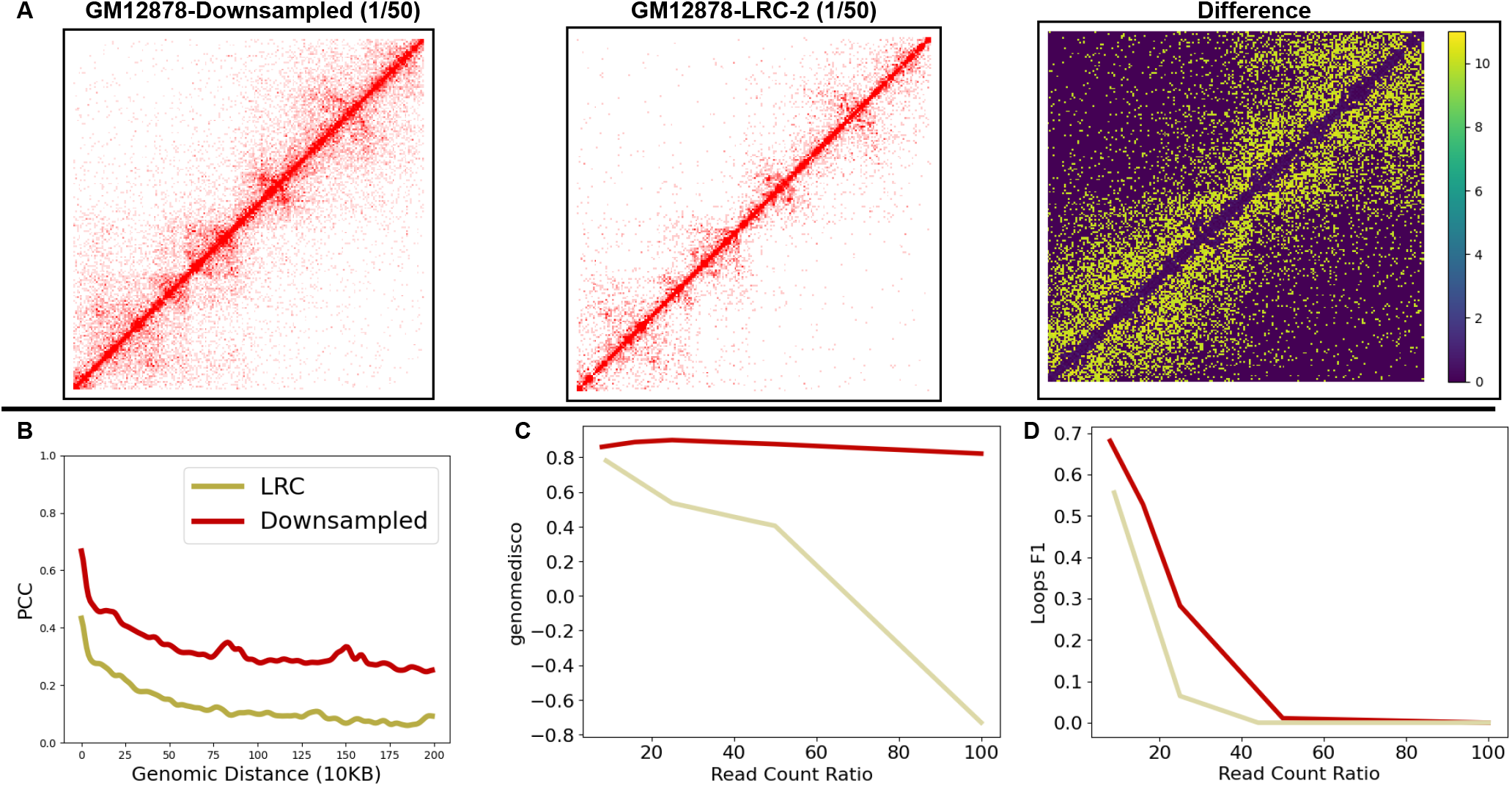
Comparing the distributions of downsampled Hi-C datasets with the experimentally generated LRC datasets. **A** Shows a pixel-wise comparison of 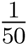 experimentally generated LRC dataset with a similarly downsampled dataset for GM12878 chromosome 22 region 41-43 Mbp. As we move further from the diagonal, the difference amplifies (signified by yellow). **B** We compare Pearson’s Correlation (on the y-axis) of the LRC dataset (beige) and Downsampled dataset (red) with 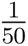 sparsity against the HRC dataset for strata with increasing genomic distances (on the x-axis). The correlation is always smaller for experimentally generated LRC datasets across all genomic distances, and the difference increases as we move further away from the diagonal. **C** Compares GenomeDISCO score of experimentally generated LRC datasets against a similarly sparse downsampled dataset. Our results show that the GenomeDISCO score is lower for experimentally generated LRC datasets compared to an equally sparse downsampled dataset. We compare the Biological Features recovery F1 score against the HRC feature set for LRC and Downsampled datasets on the GM12878 cell line. F1 scores decay more steeply for experimentally generated LRC datasets.

**Figure 5:**
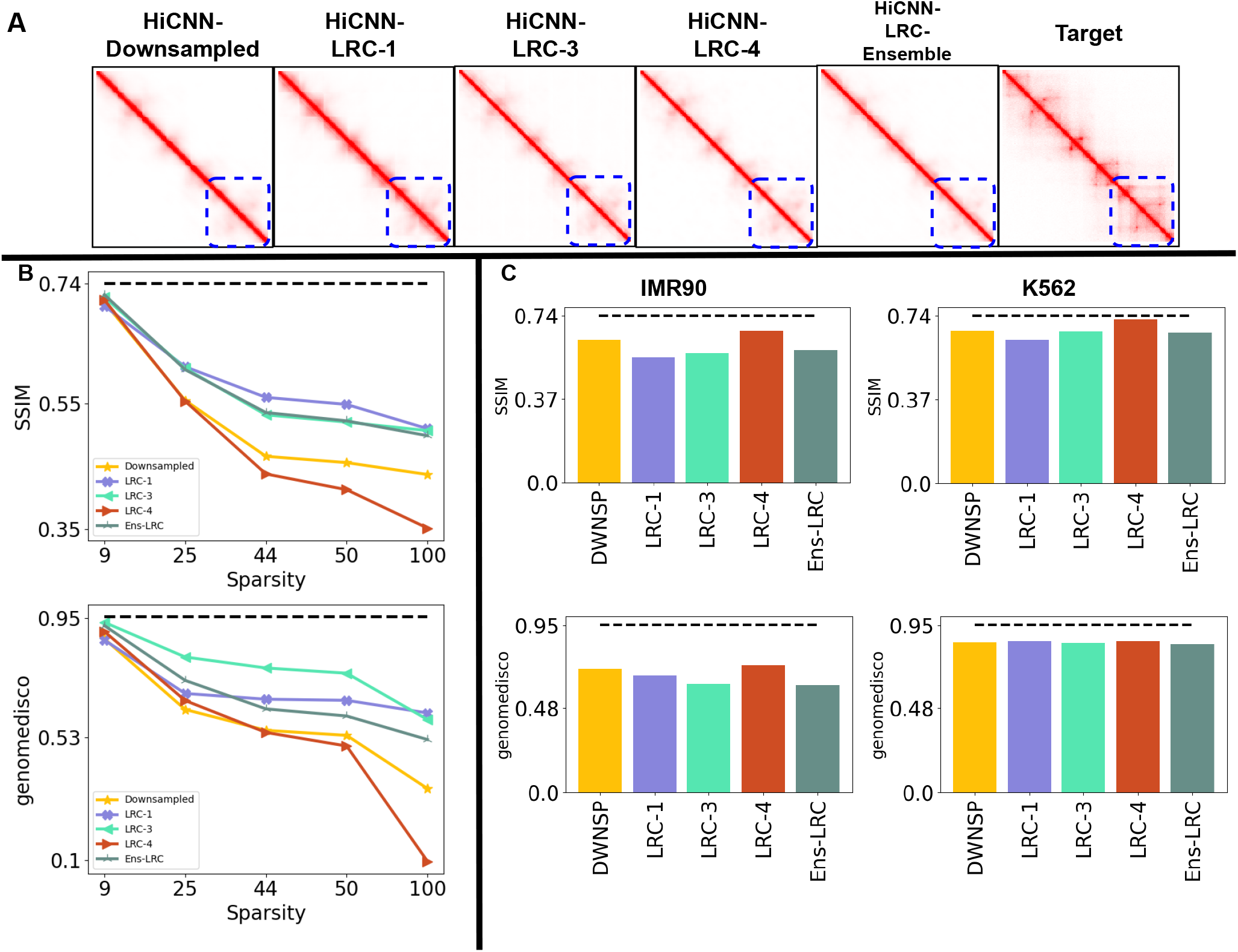
Retraining HICNN with experimentally generated LRC datasets. **A** We visualize the outputs of our methods on GM12878-LRC-2’s with 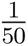 on region chr22:41-43Mbp and we show that. **B** We quantify the decrease in performance of upscaling methods as we increase the sparsity (on the x-axis) of GM12878 LRC datasets on SSIM and GenomeDISCO metrics (on the y-axis). We observe that retraining with LRC datasets or an ensemble of LRC datasets improves performance, with LRC-3 showing the most improvement in the GenomeDISCO metric and retraining with LRC-1 providing the most improvement for the SSIM score. All versions struggle to achieve scores similar to the biological replicate shown as the dotted black line. **C** We compare the performance of these methods for both IMR90 and K562’s LRC datasets. All methods perform similarly, with HiCNN trained with downsampled datasets performing better on the IMR90 dataset.

We further investigate the impact of this distributional shift by comparing the content and biological information of downsampled datasets against experimentally generated LRC datasets. Figure 4 **C** shows a steeper degradation in GenomeDISCO scores in LRC matrices compared to the downsampled matrices. Given that GenomeDISCO compares the underlying structural topology rather than bin-wise similarity, our results in Figure 4 **C** show there is a steeper loss in structural information in LRC Hi-C datasets compared to downsampled datasets. Lastly, we quantify the impact of this loss of structural information by comparing the chromatin loop F1 scores between the downsampled and experimentally generated LRC Hi-C contact maps. Unsurprisingly, Figure 4 **D** shows chromatin loop F1 scores decay more sharply for LRC datasets, while F1 scores approach 0 for both datasets. Our findings across all datasets, as shown in the Supplementary Figures S5 and S6 on other metrics and biological features, further support a more significant information loss in experimentally generated LRC datasets that scales with the sparseness of data. In the cross-cell-type setting, we find an interesting trend that suggests IMR90-LRC-1 has more loss in structure information than K562-LRC-1 even though it has higher sparsity suggesting the existence of cell-specific experimental artifacts. Our analysis identifies the distributional shift between the downsampled and LRC datasets as a crucial component that leads to poor generalizability of deep-learning based models. We further show that upscaling LRC datasets is substantially more complicated due to a lack of structure in LRC Hi-C matrices and their lower similarity to the target HRC Hi-C matrices.

**Figure 6:**
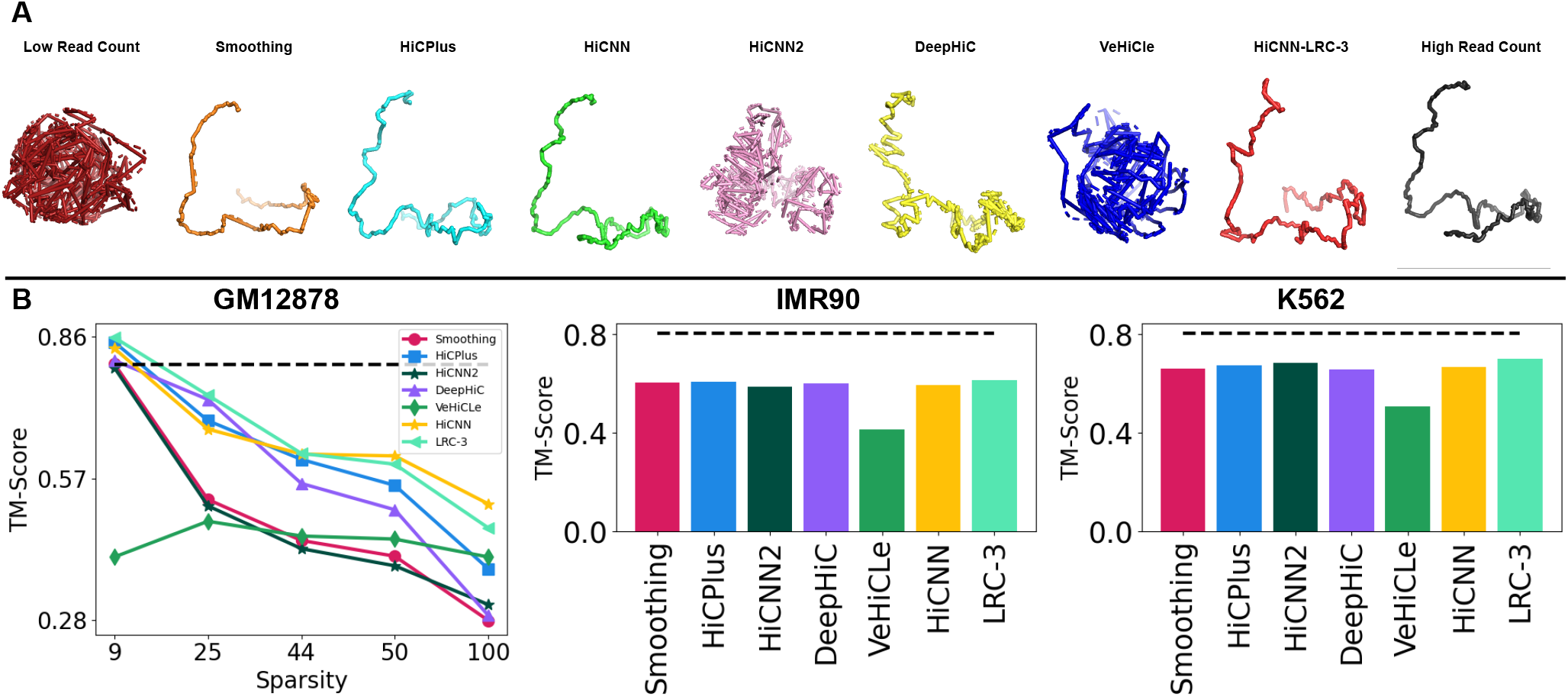
3D reconstruction of Chromosome 22:41-43Mbp region. **A**Most models including HiCNN-LRC-3 produce highly similar 3D structures of the chromatin. **B** Our Quantitative analysis on 3D reconsturction score comparison suggests retraining improves performance in certain cases while performs similarly to HiCNN on average

### 3.4 Retraining deep learning models with experimentally generated datasets improves performance

Our results in the previous sections show that the distribution of experimentally generated LRC Hi-C data differs from the downsampled Hi-C datasets. Given this insight, we pick the best model - HiCNN - and retrain it with an experimentally generated GM12878 LRC datasets. The core idea behind retraining is to expose the deep-learning model to a more realistic data distribution during the training phase to improve generalizability in the testing phase. We retrain the HiCNN model on each LRC dataset individually (LRC-1 – 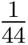 reads of HRC data, LRC-3 – 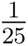 reads of HRC data, and LRC-4 –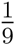 reads of HRC data). We also train on an ensemble of aforementioned LRC datasets (Ensemble-LRC) as a means to expose the model to a range of realistic distribution of Hi-C datasets during the training phase. Figure 5 **A** shows that even with retraining, HiCNN is still unable to recover the sub-TADs in the TAD cluster highlighted with a dotted blue square even when retrained with an experimentally generated LRC dataset. Figure 5 **B** shows the SSIM and GenomeDISCO scores of three versions of HiCNN trained with LRC-1, LRC-3, or LRC-4 GM12878 datasets and two versions of HiCNN trained either with an ensemble of LRC datasets or ensemble of downsampled datasets. We evaluate the performance of these models on five GM12878 datasets with varying levels of read count sparsity. We find that the versions of HiCNN retrained with LRC datasets with high sparsity (LRC-1 and LRC-3 and Ensemble-LRC) improve the deep learning model performance on average by 11%, 10%, and 10% on SSIM and 18%, 31%, and 16% on GenomeDISCO in comparison to HiCNN trained with downsampled datasets. Conversely, we observe a reduction in performance when retrained with a low sparsity (LRC-4) on average by 6% on SSIM and 8% on GenomeDISCO compared to when trained with downsampled datasets. However, we observe the most performance improvement of 16%, 16%, 14% on for LRC-1, LRC-3 and Ensemble-LRC, respectively on SSIM. We also see improvement of 75%, 68%, 48% for LRC-1, LRC-3, LRC-4 and Ensemble-LRC, respectively on the GenomeDISCO metric when upscaling GM12878-LRC-5, which is the sparsest dataset having 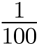 reads.

Next, we use IMR90-LRC-1 and K562-LRC-1 datasets to compare the performance of HiCNN retrained with experimentally generated LRC datasets against HiCNN trained with downsampled datasets in the cross-cell type setting. Figure 5 **C** shows that when evaluated on K562-LRC-1 dataset, the retrained versions of HiCNN performed similarly to the original HiCNN on both SSIM and GenomeDISCO. Whereas, we observe a reduction of 8%, 9%, and 7% when trained with LRC-1, LRC-3, and Ensemble-LRC and conversely an increase of 7% when trained with LRC-4 on SSIM scores, respectively. We observe similar GenomeDISCO scores across all variants of HiCNN when evaluated on IMR90-LRC-1 dataset. Retrained HiCNN does not generalize to unseen cell-types. We hypothesize this happens because distributional differences across cell-type are independent of the read counts. Overall, we demonstrate that retraining HiCNN using LRC datasets is a promising approach towards improving the generalizability of deep-learning based models because it exposes them to a more realistic Hi-C contact map distribution during the training phase.

### 3.5 Downstream analysis to quantify the utility of upsampled Hi-C contact maps

To determine whether the upscaled Hi-C matrices retain necessary biological signals, we analyze the 3D structure of chromatin and biological features. HiCNN, HiCPlus, and DeepHiC [8, 15, 16, 29] presented these results for upscaled outputs from synthetically downsampled Hi-C maps. We perform this downstream analysis on upscaled experimentally generated LRC datasets to provide insights into the performance for real-world applications.

#### 3.5.1 Retrained deep learning models generate highly accurate 3D structure of chromatin

For 3D chromatin reconstruction analysis, we use the upscaled Hi-C maps produced by the all the methods from all seven experimentally generated LRC input matrices on GM12878, IMR90, and K562. For this, we incorporate 3DMax [20] into the Hi-CY framework, which generates the 3D structure for the 200*×*200 Hi-C sub-matrices for all of the sub-matrices in the test chromosomes. Next, we compare the spatial conformation of the chromatin using the TM-score, where a higher TM-score is proportional to a higher structural similarity of the chromatin. Our analysis follows the VeHiCle paper[7], where the recovered 3D structure from HRC Hi-C contact map is compared with the 3D structure recovered from the upscaled Hi-C matrices to show that the upscaled matrices have the same structure. We also visualize 3D models of the region chr22:41-43Mbp for upscaled GM12878 in Figure 6 **A**, which shows that models except for HiCNN2 and VeHiCle are similar in their overall 3D topology compared to the 3D model generated with HRC Hi-C contact map.

We summarize the TM-Scores in Figure 6 **B**. We observe that, on average, both HiCNN-Baseline and HiCNN-LRC-3 show similar performance on 3D reconstruction analysis with scores within 1% of each other. However, although the gains are marginal, we observe that HiCNN-LRC-3 outperforms all methods on both K562 and IMR90 cell lines. Our results suggest retraining improves 3D reconstruction performance or performs similarly to the baseline version of HiCNN, highlighting the value of retraining for recovering biologically informative Hi-C matrices.

#### 3.5.2 Deep learning models have similar performance for recovering biological features

We incorporate Chromosight [18] into the Hi-CY evaluation pipeline to recover and compare biological features, including chromatin loops, TADs, and DNA-hairpins. To compare the utility of the upscaled Hi-C matrices in recovering biological features, we detect features on all seven experimentally generated LRC datasets upscaled from the complete set of models, including HiCNN retrained with the LRC-3 dataset. In Figure 7, we compare the F1 scores across all seven datasets for all three features we recover. For all models, we observe a trend similar to GenomeDISCO and SSIM, that F1 decreases as we increase the sparsity of the LRC dataset. Retraining with the LRC-3 dataset helps recover features far from the diagonal such as chromatin loops and TADs, particularly in IMR90, K562, and lower sparsity GM12878 cases. Regardless, the performance improvements, as observed earlier, are marginal. Surprisingly, HiCNN2 outperforms other methods in the loops and DNA-Hairpin case in feature recovery analysis on GM12878 LRC datasets. This provides evidence of a need for multifaceted research that explores results across multiple scenarios and metrics to develop a holistic understanding of the costs as well as benefits of using deep-learning based models. Currently, all existing methods struggle to recover biological features accurately, particularly in a cross-cell type setting and on sparse Hi-C datasets, as we see a steep degradation in F1 scores in both scenarios. We observe similar scores or negligible improvements even when we retrain HiCNN with an experimentally generated LRC dataset. This suggests a need to revisit architectural assumptions as well, including formulating Hi-C as a 2D image when the underlying measurement data corresponds to a 3D structure.

**Figure 7:**
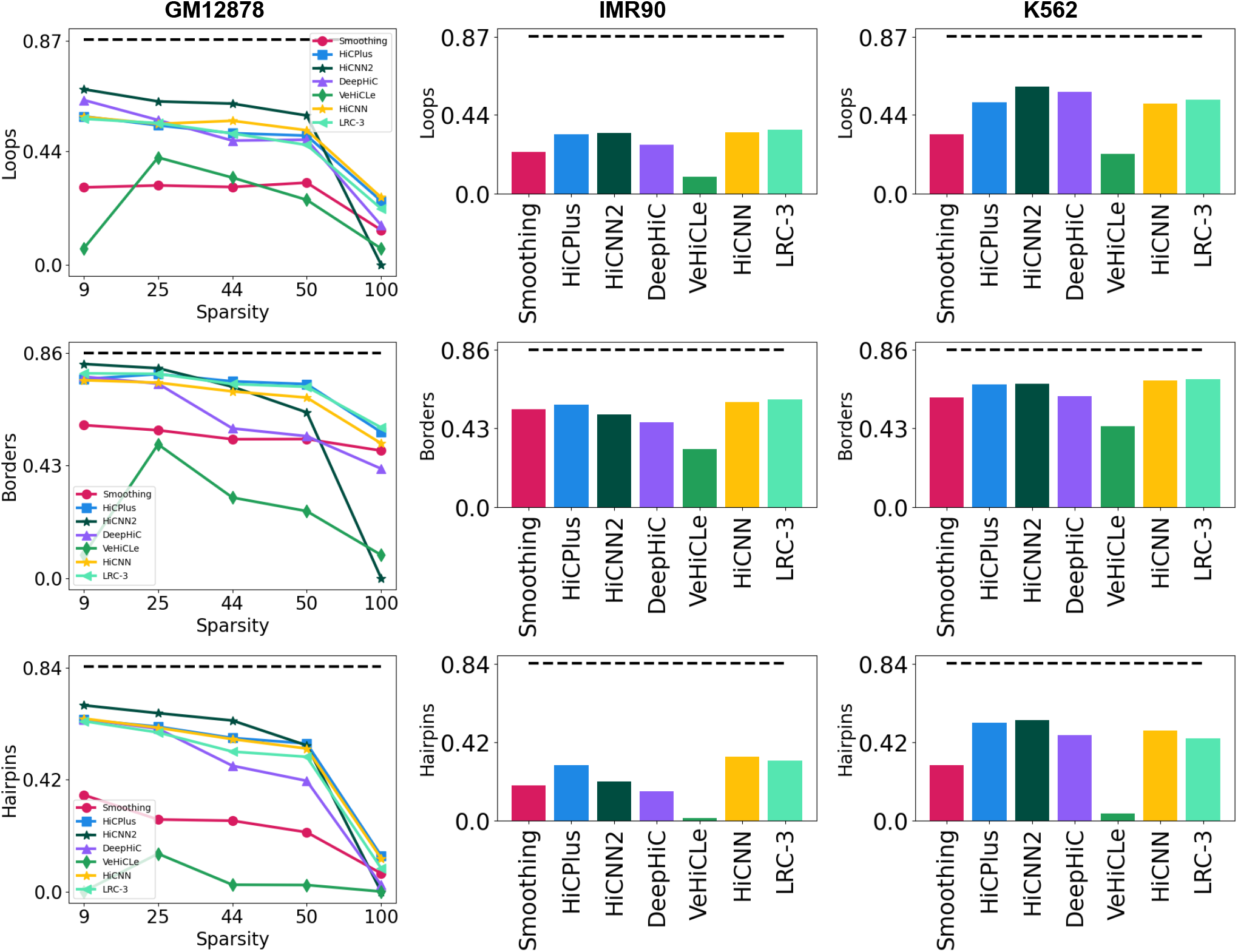
Biological feature comparison. Our Quantitative analysis by comparing F1 scores, computed by comparing the recovered feature set with the HRC feature set suggests across Chromatin Loops, TADs, and DNA-Hairpins across all three cell lines for all of the methods suggest that all methods, including retrained HiCNN, struggle to recover meaningful biological features in highly sparse settings. While retraining improves performance, the improvements are marginal similar to the previous analysis.

## 4 Conclusion

We develop the Hi-CY framework that explores how deep-learning based Hi-C upscaling methods perform in their intended scenarios which is to upscale experimentally generated sparse Hi-C datasets. Our results strongly suggest that existing deep-learning based methods do not generalize to their real-world use case. To provide potential solutions to improve generalizability, we explore retraining with dataset augmentations (such as adding Gaussian noise to the training datasets show in the Supplementary Figure S8), ensembling of multiple downsampled datasets, and training with experimentally generated Hi-C datasets. Our analysis found that retraining with experimentally generated sparse datasets was the most promising approach to improving generalizability. Although we did not observe a significant improvement in 3D structure generation or recovering chromatin loops, TADs, and DNA-hairpins analysis tasks, we still recommend retraining with experimentally generated Hi-C datasets to improve their generalizability beyond the training datasets. Our Hi-CY framework provides a simple yet robust pipeline to streamline training and evaluation of Hi-C upscaling methods with real-world Hi-C datasets on correlation, HiC-similarity, and downstream analysis-based metrics.

Another trend we observe in our evaluation is that among the currently available techniques, HiCNN, which uses a non-adversarial model, outperforms the GAN-based models in generalizing to sparse experimentally generated Hi-C datasets. Interestingly, we observe that an even simpler method HiCPlus (3-layer CNN), performs comparably to HiCNN (a 54-layer CNN) on most metrics across most datasets. In future work, we plan to investigate if this similar performance stems from formulating Hi-C data as an image and the upscaling as a super-resolution task. We plan to explore performance gains from using graph-based architectures to capture the underlying geometry of the DNA molecule more accurately. In conclusion, we present the Hi-CY framework that provides a comprehensive evaluation toolkit coupled with real-world datasets that supports the development of future models by investigating their performance on intended use cases.

## Acknowledgments

We thank Justin Zook from the National Institute of Standards and Technology (NIST) for providing incredibly valuable feedback on the draft. We also would like to thank Hyeyeon Hwang for sharing her valuable experience about Hi-C downstream analysis methods and having many helpful discussions with us on the draft. Lastly, we would also thank the Compbio Core at brown to for their support in setting up the resources required to conduct experiments.

## Disclaimer

Certain commercial equipment, instruments or materials are identified to adequately specify experimental conditions or reported results. Such identification does not imply recommendation or endorsement by NIST, nor does it imply that the equipment, instruments or materials identified are necessarily the best available for the purpose.

## Investigating the performance of deep learning methods for Hi-C resolution improvement

### Supplementary Material

#### S1 Retraining of HiCNN, HiCNN2 and HiCPlus

We retained the original implementation of these methods wherever we could. We changed the output objective from predicting raw counts to predicting a matrix with values between 0-1, similar to DeepHiC, and for these three, we replaced the SGD optimizer with the Adam optimizer to stabilize the training process and improve the performance. In the figure S1 we show the training curve for these methods for all the four version we retrained.

**Figure S1:**
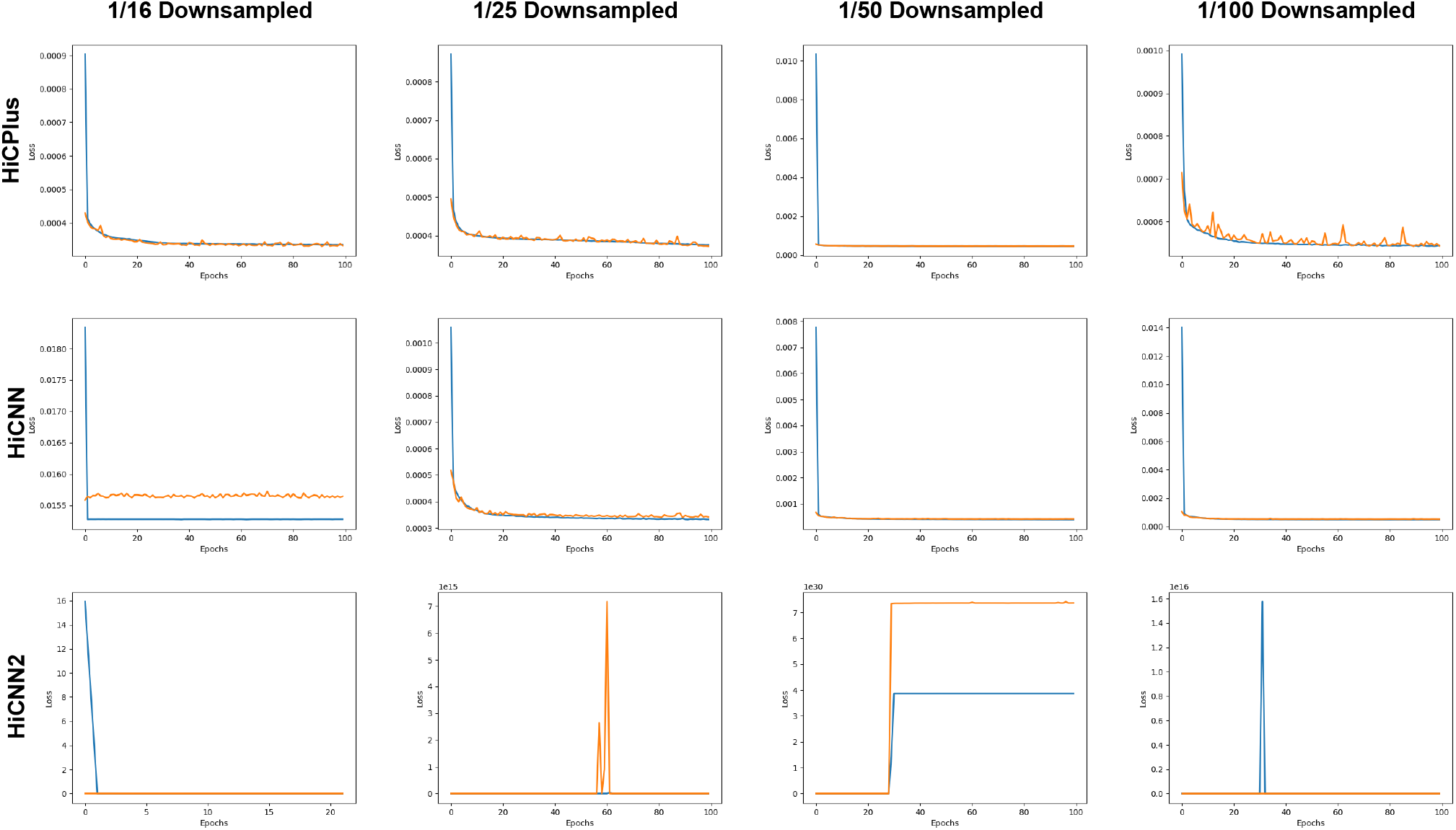
This figure shows the loss curves for the HiCNN, HiCNN2 and HiCPlus models on four version we trained with each downsampling ratio. We use the version of model (across all the epochs) that minimizes the validation loss.

#### S2 Details of the evaluated methods

- **Gaussian Smoothing** We applied the Gaussian Smoothing filter, commonly used as a baseline [29], to establish emphasize the performance benefits of the deep-learning-based methods. This method uses a 2D kernel of shape *n × n* where n is a hyper-parameter. Each kernel value follows a 2D Gaussian distribution with hyperparameters *σ_x_* and *σ_y_* that represent the relative importance of neighboring features in prediction. The smoothing operation convolves this kernel on each pixel of a 2D image, or in our case, a HiC matrix read count, to update its value. This updated value contains the average of the neighboring values weighted by 2D Gaussian distribution in the kernel. This smoothing operation removes noise in the input matrix and improves the peak signal to noise ratio (PSNR) at the cost of blurring the features. For our experiments, we performed a grid search and found the kernel size of *n* = 17 and *σ_x_* = *σ_y_* = 7 to give the best reproducibility score on the validation set of LRC HiC matrices.
- **HiCPlus** [29] HiCPlus is the first application of deep learning to improve HiC resolution. It utilizes a standard three-layer convolutional neural network (CNN) architecture to upscale a low-resolution HiC matrix by mapping it to the target high-resolution matrix. To optimize its parameters, HiCPlus uses a mean squared error (MSE) loss. HiCPlus inputs HiC matrices as sub-matrices of size 40*×*40 binned at 10Kbp resolution and taken from 2 Mbp distance from the diagonal. All the following deep learning-based models follow the same input formulation. For this study, we make certain modifications to the original code to make HiCPlus comparable to the more recent implementations. For example, the original HiCPlus was trained to generate raw read counts rather than normalized HiC matrices. Therefore, we retrain the model to work with normalized high-resolution and low-resolution pairs of HiC sub-matrices. Moreover, the original implementation was trained to upscale only the matrices that had been downsampled to 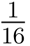 of the read counts of a high-resolution HiC map. We retrain the HiCPlus with three additional input HiC datasets with 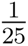, 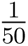, and 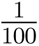 downsampling ratios to explore the performance on matrices beyond the original downsampled version.
- **HiCNN** [16] HiCNN model also uses a CNN architecture like HiCPlus, except that it consists of a much deeper 54-layer neural network with ResNet layers [6]. The choice of ResNet layers provides two key benefits - (1) it provides additional architectural complexity to learn relevant non-linear relationships between the inputs and the outputs, and (2) they train significantly faster than regular CNN layers saving substantial time during training. In addition, the skip connections in ResNet layers further avoid model overfitting on the data. HiCNN, similar to HiCPlus, uses an MSE loss to optimize its parameters and learn the mapping between the input low-resolution matrix and the high-resolution target matrix. It also produces raw read counts and is trained with downsampling ratios of 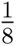 and 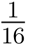 and 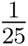. Therefore, for consistency, we retrain the model with additional datasets with downsampling rations of 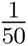 and 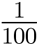 and standardized values.
- **HiCNN2** [17] HiCNN2 extends the architecture of HiCNN by ensembling multiple methods to achieve better resolution. There are three different ConvNets in HiCNN2; the first ConvNet is similar to HiCNN in terms of how it uses global and local residual learning; it also concatenates features across residual blocks to improve performance. The second ConvNet is a modified version of VDSR [12] which only uses the global residual learning. The third ConvNet is the HiCPlus model. Each model produces a 28×28 output matrix that is then combined through weights tuned during the training process. This method, similar to HiCPlus and HiCNN, also predicts raw read counts and is only trained with 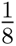 and 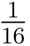 and 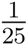 downsampling ratios. Similar to both HiCNN and HiCPlus, we retrain HiCNN2 to predict normalized read counts across all four downsampling ratios.
- **HiCGAN** [15] HiCGAN paper argues that the Mean Squared Loss function used in both HiCNN and HiCPlus causes these models to generate over-smooth matrices. Therefore, it proposes using a Generative Adversarial Network (GAN) model for the HiC resolution improvement task. A GAN architecture consists of (1) a generator and (2) a discriminator. The generator’s objective is to produce data that increasingly resembles the original distribution, and the goal of the discriminator is to identify fake (generated) data from the original data. This coupled training causes both models to get iteratively better at their tasks. HiCGAN uses a specialized form of GAN, which is called the conditional GAN (cGAN). In cGANs the generator produces an output conditional on the provided input instead of a random noise input. To optimize the parameters in the model, HiCGAN uses the discriminator loss and the MSE loss to generate matrices that are highly similar to the target HiC matrices. The HiCGAN paper shows that this method produces better-quality HiC matrices with sharper and more prominent features than the previous method.
- **DeepHiC** [8] DeepHiC paper, like HiCGAN [15], argues that the Mean Squared Loss function used in both HiCNN and HiCPlus causes these models to generate over-smooth matrices. Therefore, it uses Generative Adversarial Network (GAN) model for the HiC resolution improvement task. A GAN architecture consists of (1) a generator and (2) a discriminator. The generator’s objective is to produce data that increasingly resembles the original distribution, and the goal of the discriminator is to identify fake (generated) data from the original data. This coupled training causes both models to get iteratively better at their tasks. DeepHiC substantially revises the previously proposed loss functions to contain additional functions that include Total Variation Loss and Perceptual Loss. These loss function along side Mean Squared Error Loss and Discriminator Loss causes the model to generate matrices with sharper features that are more biologically informative [8]. The paper also shows that training the deep learning models on standardized HiC matrices improves their performance further. Moreover, the paper trains the DeepHiC model with a downsampling ratio of up to 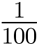 to have model weights available for even the sparsest real-world HiC matrices.
- **VeHiCLE** [7] VeHiCLE is another GAN-based model like HiCGAN and DeepHiC. However, it makes additions to both the model architecture and loss functions used while training. Apart from using a GAN architecture, it also trains a variational auto-encoder (VAE). The output obtained by passing the HiC matrices through the trained VAE is used in a loss function to train the GAN. This loss obtained from the VAE is called the variation loss. VeHiCLE also uses the adversarial and MSE losses seen in previous methods. However, it introduces yet another loss called insulation loss. The insulation loss is a biologically inspired loss that utilizes insulation scores used to identify TADs in a HiC contact matrix. VeHiCLE is trained on input and target matrices of sizes 269*×*269, unlike the previous methods that used 40*×*40 sub-matrices as inputs. The paper shows that this increased matrix size improves performance, thus hypothesizing that the 40 *×* 40 matrices are too small to adequately capture information about large-scale HiC features (such as TADs). We created new datasets for VeHiCle that had HiC sub-matrices of size 269×269 to ensure a fair comparison of VeHiCle with other deep learning based methods.

#### S3 Details of the evaluation metrics

- **Structural Similarity Index Measure (SSIM)** Structural Similarity Index Measure (SSIM) is a metric that measures the perceived perceptual quality of an image against an original undistorted and higher quality image. SSIM measures this perceptual quality by comparing the luminance, contrast, and structural properties in small local regions of the images. A weighted sum of these properties allows SSIM to assign a similarity score that closely mimics the way humans perceive differences in images. However, we postulate that the HiC contact maps should be compared based on their underlying biological properties instead of their visual similarities. Therefore, assigning a similarity score based on SSIM score may hold little biological relevance and might lead to misleading conclusions about the quality of the generated datasets.
- **Pearson Correlation Coefficient (PCC)** Pearsons Correlation Coefficient (PCC) is a linear measure of the correlation between two sets of data distribution. PCC is a ratio of covariance between two variables and its product with their standard distribution. This metric essentially measures covariance between the two datasets, normalized to have a value between −1 and 1. Here, −1 or 1 values imply highly negatively or positively correlated, respectively, and a 0 value implies no correlation between the two datasets.
- **Spearman’s rank Correlation Coefficient (SCC)** Spearman’s rank Correlation Coefficient (SCC) measures the statistical dependence between the rank of two variables. This measure essentially captures how well two variables can be described using a monotonic function. SCC between two variables is equal to the PCC of the rank of variables. Thus, SCC has a value of +1 or −1 when either of the variables is a perfect monotone of the other. It has a value of 0 when they do not correlate monotonically.
- **Hi-C-Rep** [26] Hi-C-Rep measures the reproducibility between two Hi-C datasets based on spatial features such as distance dependence and domain structure. To enhance the domain structure in the Hi-C matrices, Hi-C-Rep applies a mean filter, which filters out the stochastic noise that can potentially arise from the experimental protocols and possibly also through low read counts in the dataset. Hi-C-Rep then stratifies the read counts in the matrix based on their distance and measures the correlation between each stratum. The correlations between each stratum are then combined using a weighted average of each stratum, where the weighting coefficients are calculated using the Cochran-Mantel-Haenszel (CMH) statistic. Hi-C-Rep reports a score between −1 and 1, similar to Pearsons’s Correlation and Spearman’s, with scores close to 1 representing high similarities between the Hi-C matrices.
- **GenomeDISCO** [24] The GenomeDISCO method focuses its correlation analysis on the property that high-level order structures (loops and compartments) at multiple scales between two Hi-C contact maps are similar if they are highly correlated. To leverage that, GenomeDISCO measures the correlation over a range of genomic scales by smoothing the Hi-C matrices at different intensities. GenomeDISCO uses Random Graph walks to smooth out the matrices by formulating the Hi-C data as a Graph. In this contact matrix graph, each node represents a genomic region and, edge weights represent the contact value between these regions. Random Graph walk measures the probability of finding a path between any given node pair *i* and *j* given we can only take *t* steps, where for each step *t*, the edge chosen for the walk is dependent on edge weight. GenomeDISCO then raises the power of each value in the random walk network by power of *t* to construct a smoothed contact map. The higher the value of *t*, the higher dimensional genomic features(such as A/B compartments) the smoothed contact map summarizes. Finally, to obtain the reproducibility score, GenomeDISCO computes the area under the curve of the *t* against the L1 distances between the smoothed contact maps. Since maximum L1 distance between contact maps can be 2, the reproducibility is calculated by *score* =1*−combined−distance* which gives the score in range [*−*1,1]. Where value of 1 represents a high similarity between the input contact maps.
- **Hi-C-Spector**[25] Hi-C-Spector computes the correlation by performing a spectral analysis of the input Hi-C matrices. Hi-C-Spector’s analysis is centered around the observation that the first two eigenvectors of the Hi-C matrix correspond to the higher dimensional structures such as A/B compartments. Given that observation, two Hi-C matrices that are similar should have eigenvectors that are also similar. To compute the similarity between those eigenvectors, Hi-C-Spector does the following operations: First of all, it constructs a Laplacian Matrix *L* by subtracting the Diagonal of the contact matrix *D* from the input contact matrix *W*. In the second step, it normalizes the contact map by applying a transformation *D^−^*^1^*^/^*^2^*LD^−^*^1^*^/^*^2^. In the third step, Hi-C-Spector decomposes the normalized matrix into its eigenvectors. It keeps the first two eigenvectors and discards the rest because those vectors represent the noise in the data. In the last step, Hi-C-Spector computes a summation of pair-wise distances of the Eigenvectors of the input Hi-C contact map against the reference Hi-C map and normalizes it in the range 0-1 to assign a similarity score, where the score of 1 represents the high similarity between two contact maps.
- **QuASAR-Rep** [28] QuASAR-Rep bases its correlation analysis on the observation that in a distance matrix, as the distance between two features approaches zero, the correlation between rows of these features approaches one. QuASAR-Rep utilizes this property to compute the correlation between two Hi-C samples. It first filters out all the intra-chromosomal contacts from the input Hi-C samples and then all the rows that do not contain a non-zero contact bin within a 100 bin range of the diagonal. Next, it computes the background signal-to-distance ratio by taking the average reads of each inter-bin distance. Next, it sets up the interaction correlation matrix. The interaction correlation matrix is constructed based on all the pairwise sets of rows and columns (within a range of 100 bins of each other) in the log-transformed enrichment matrix. The values in the enrichment matrix are non-filtered counts divided by background signal to distance values. Finally, to compute the correlation between a given set of rows A and B, correlation is calculated between all the columns in a 100 bin range of both rows (excluding the filtered rows). QuASAR-Rep adds 1 to all the valid entries and takes a square root to construct the interaction matrix. The weighted interaction matrix is just an element-wise product of the correlation matrix and the interaction matrix divided by the sum of all valid interaction matrix entries. It calculates the final reproducibility score by computing the correlation between the weighted correlation matrices of the input samples. This score is 0 to 1, where one represents that Hi-C inputs are highly similar and zero represents that they are highly dissimilar.

**Figure S2:**
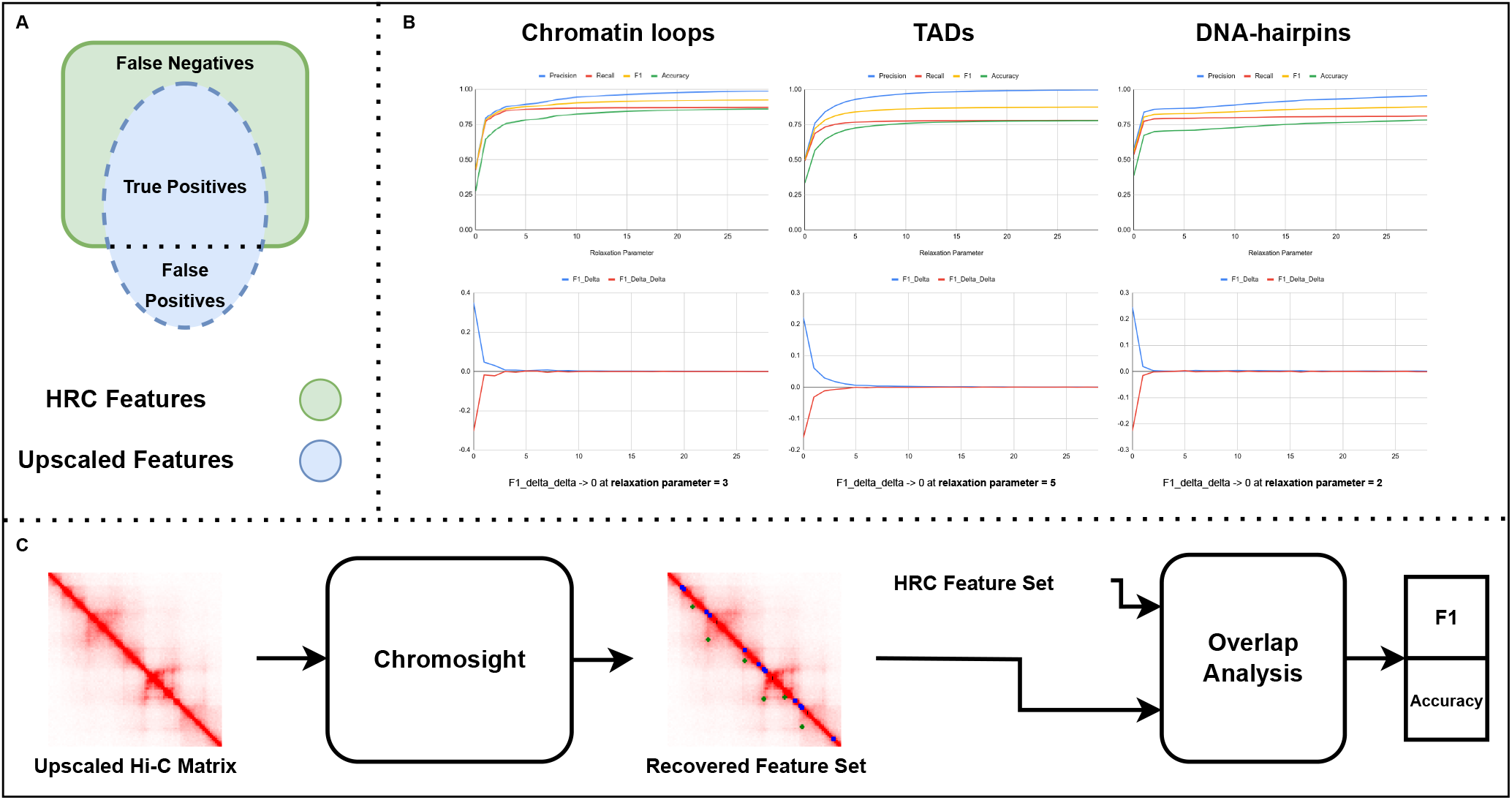
Overview of our Biological Feature Analysis Pipeline. **A** We define True Positives as features that we find to be overlapping between the upscaled Hi-C matrix and the HRC matrix. Similarly, we define features that are present in the HRC matrix and not in the upscaled Hi-C matrix as False Negatives. Conversely, features we find in upscaled matrix and not in the HRC matrix as the false postives. We compute F1 scores using these definitions. **B** We optimize the parameter “r” (relaxation paramter) for each feature, chromatin loops, TADs and DNA-hairpins. This relaxation parameter defines the overlapping radius between the position of features between the upsampled and HRC features. We tune this paramter by comparing features between the GM12878 HRC and GM12878 biological replicate cell lines and find the value of r that maximizes the F1 score while keeping the value of r small. We found a value of 3 for chromatin loops, 5 for TADs and 2 for DNA-hairpins. **C** We summarize our biological feature analysis pipeline, we first extract biological features using Chromosight and then compute F1 scores.

**Figure S3:**
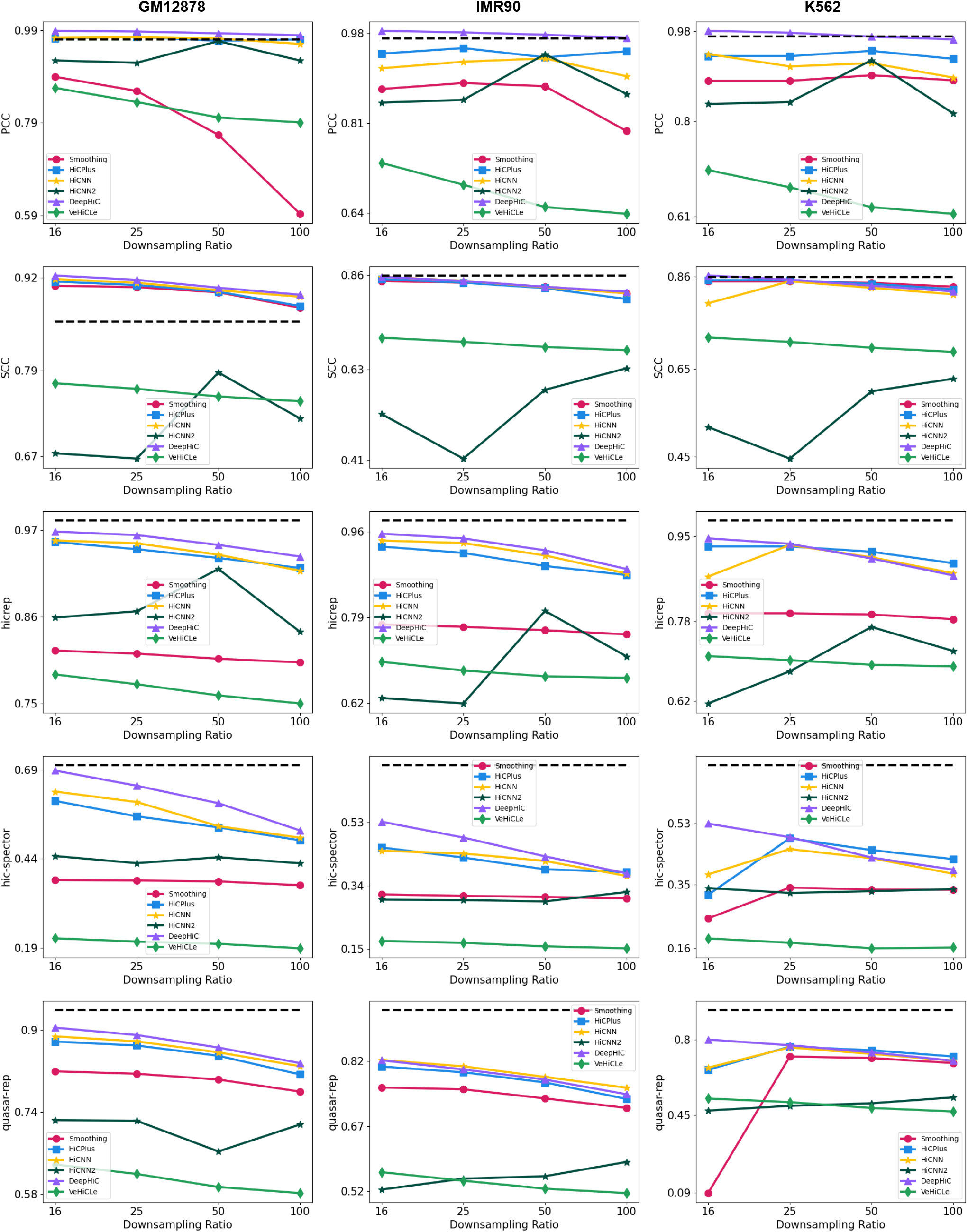
This figure shows the performance of deep-learning based Hi-C upscaling methods on computationally downsampled methods on four downsampling ratios (shown on x-axis), three cell-lines five metrics. All methods except VeHiCLe show performance very similar to the replicate shown as black dotted bold line.

**Figure S4:**
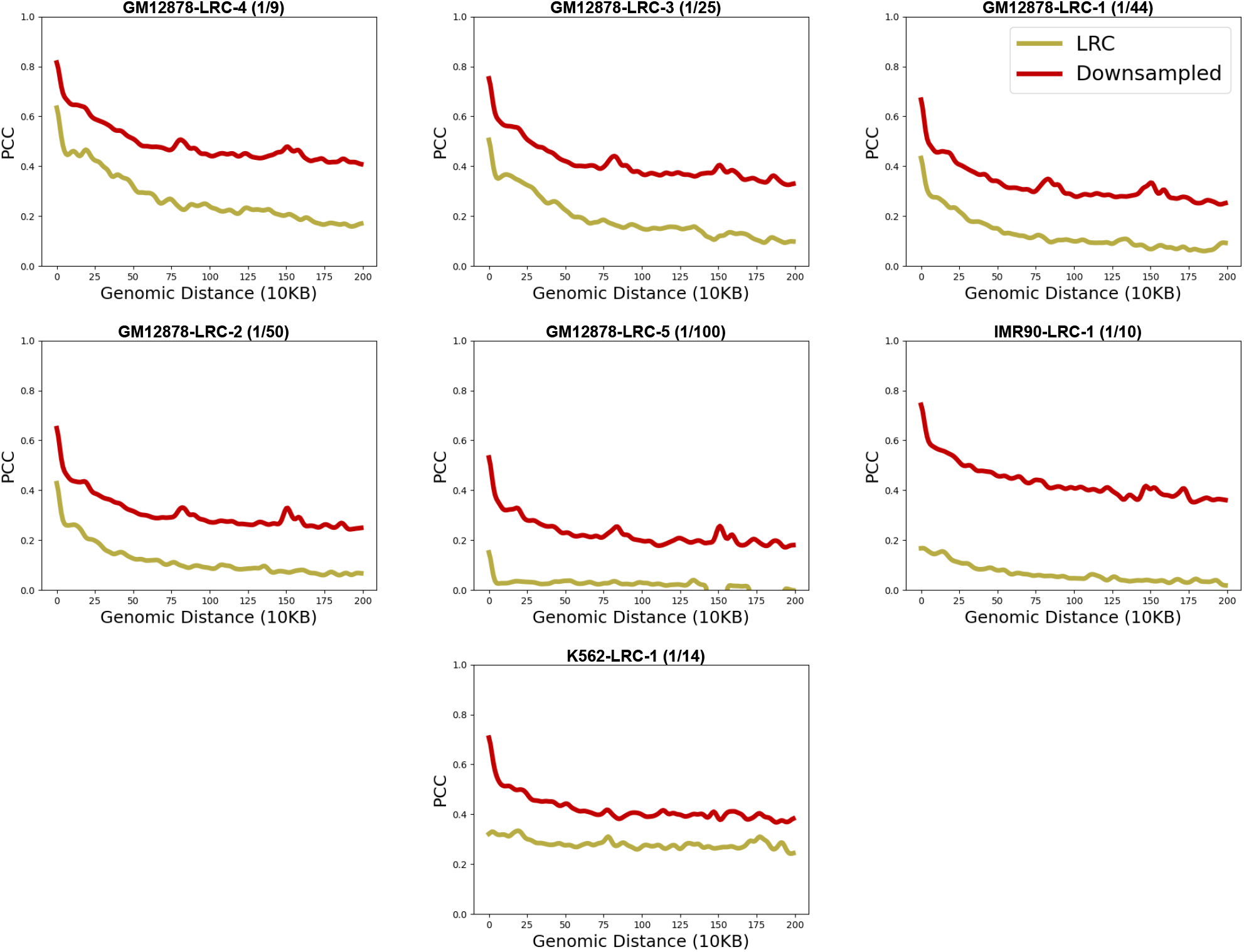
Our results on comparing the PCC (y-axis) across various genomic distances (x-axis) suggest that LRC datasets show a smaller similarity with HRC Hi-C dataset in both increasing levels of read sparsity and in cross-cell-type cases.

**Figure S5:**
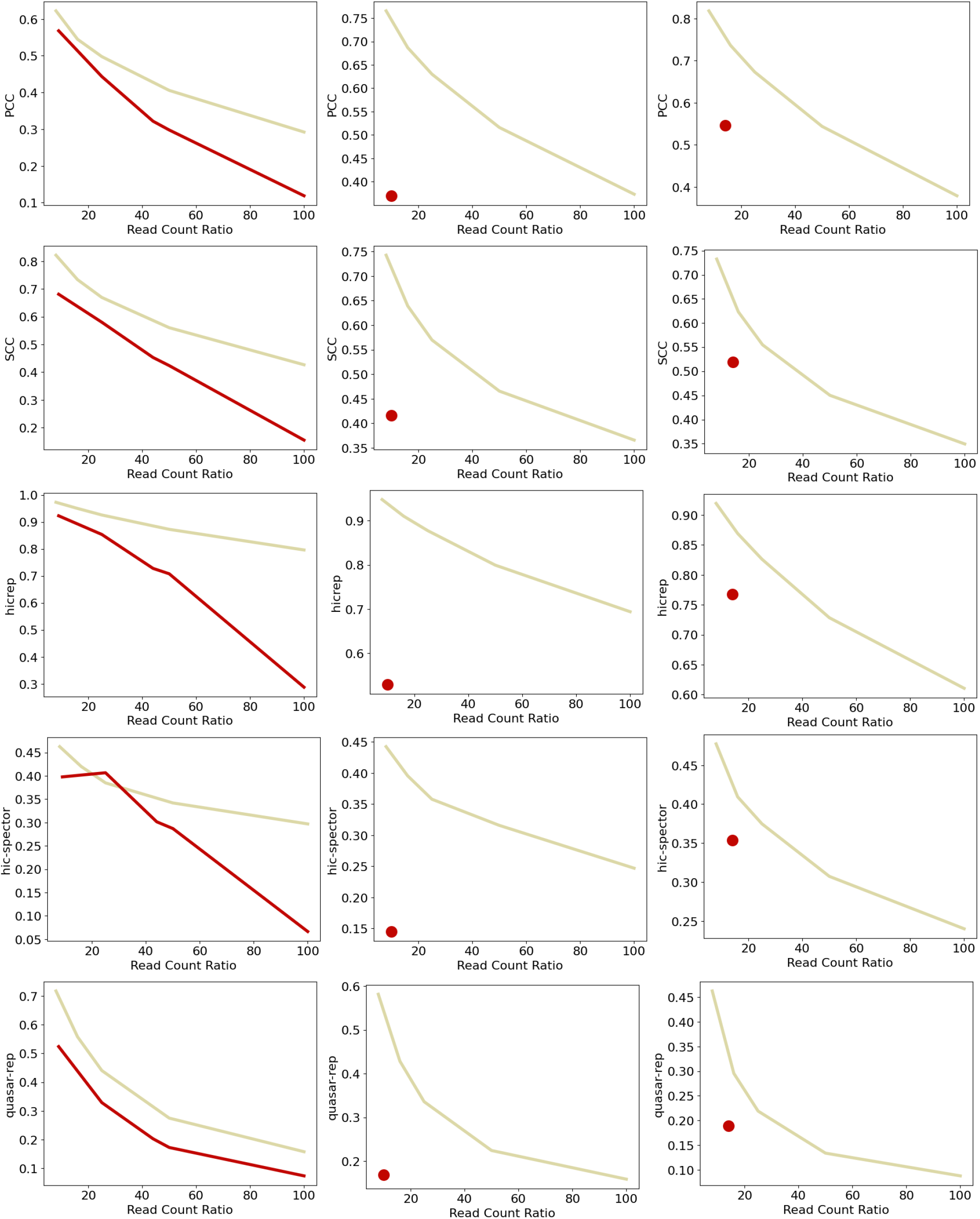
Our results on comparing the PCC (y-axis) across various genomic distances (x-axis) suggest that LRC datasets show a smaller similarity with HRC Hi-C dataset in both increasing levels of read sparsity and in cross-cell-type cases

**Figure S6:**
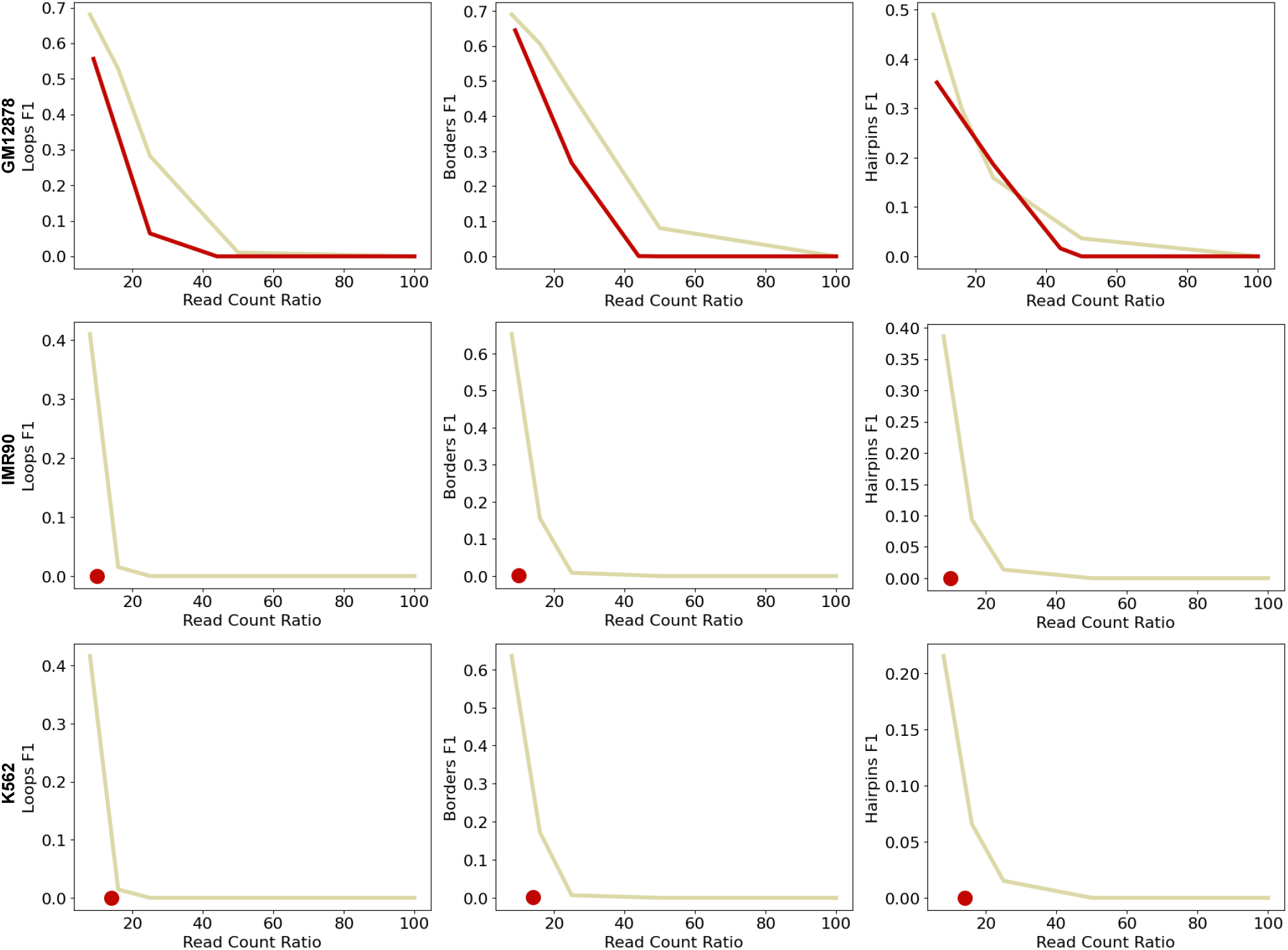
We show the impact of the distributional difference in the structural information contained in downsampled and experimentally generated LRC Hi-C contact maps by comparing scores across both correlation and Hi-C similarity metrics

**Figure S7:**
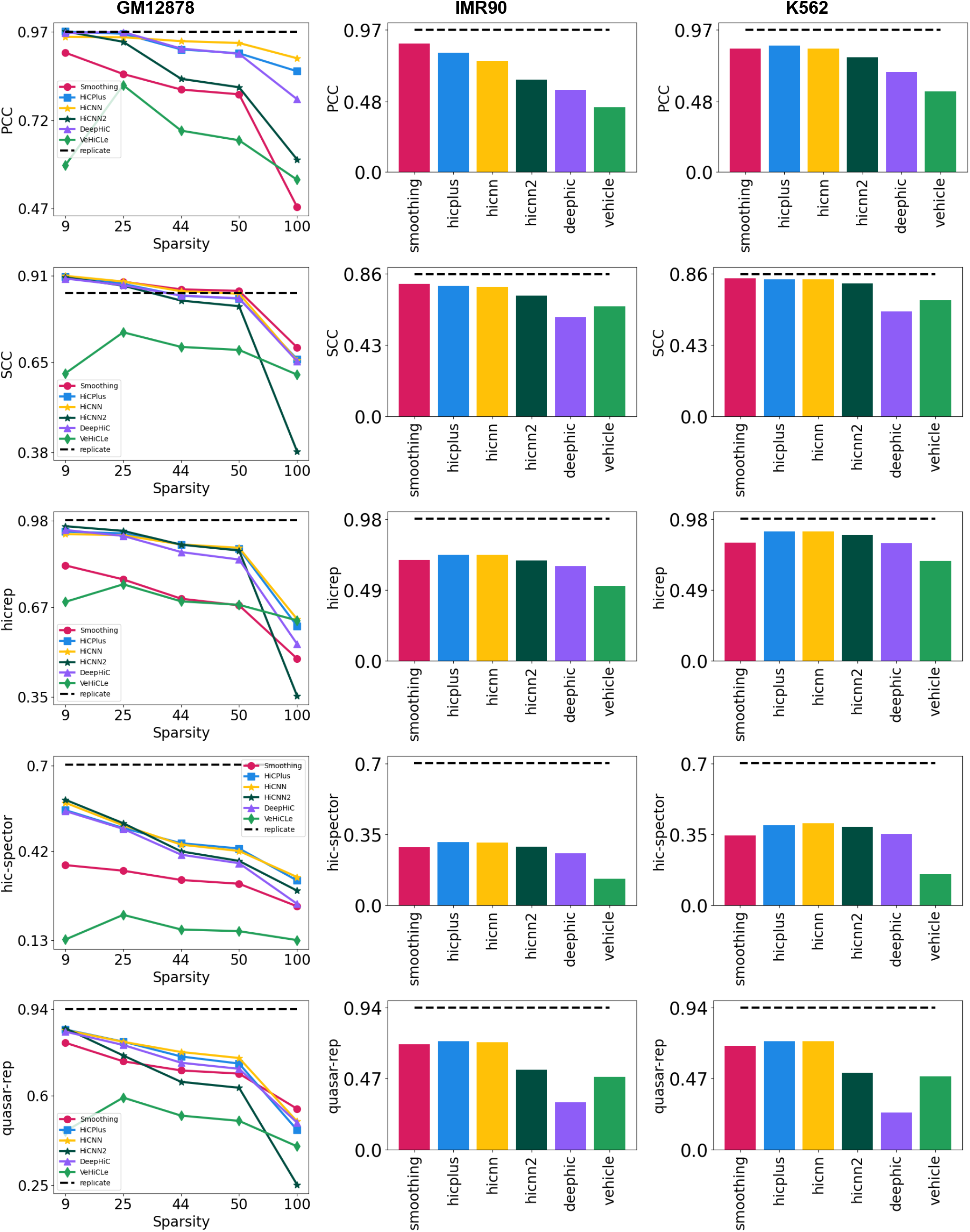
This figure shows performance on rest of the metrics, and our results highlight that there is a reduction in performance as the sparsity increases in GM12878 datasets or we evaluate these methods in a cross-cell-type (IMR90 and K562) setting.

**Figure S8:**
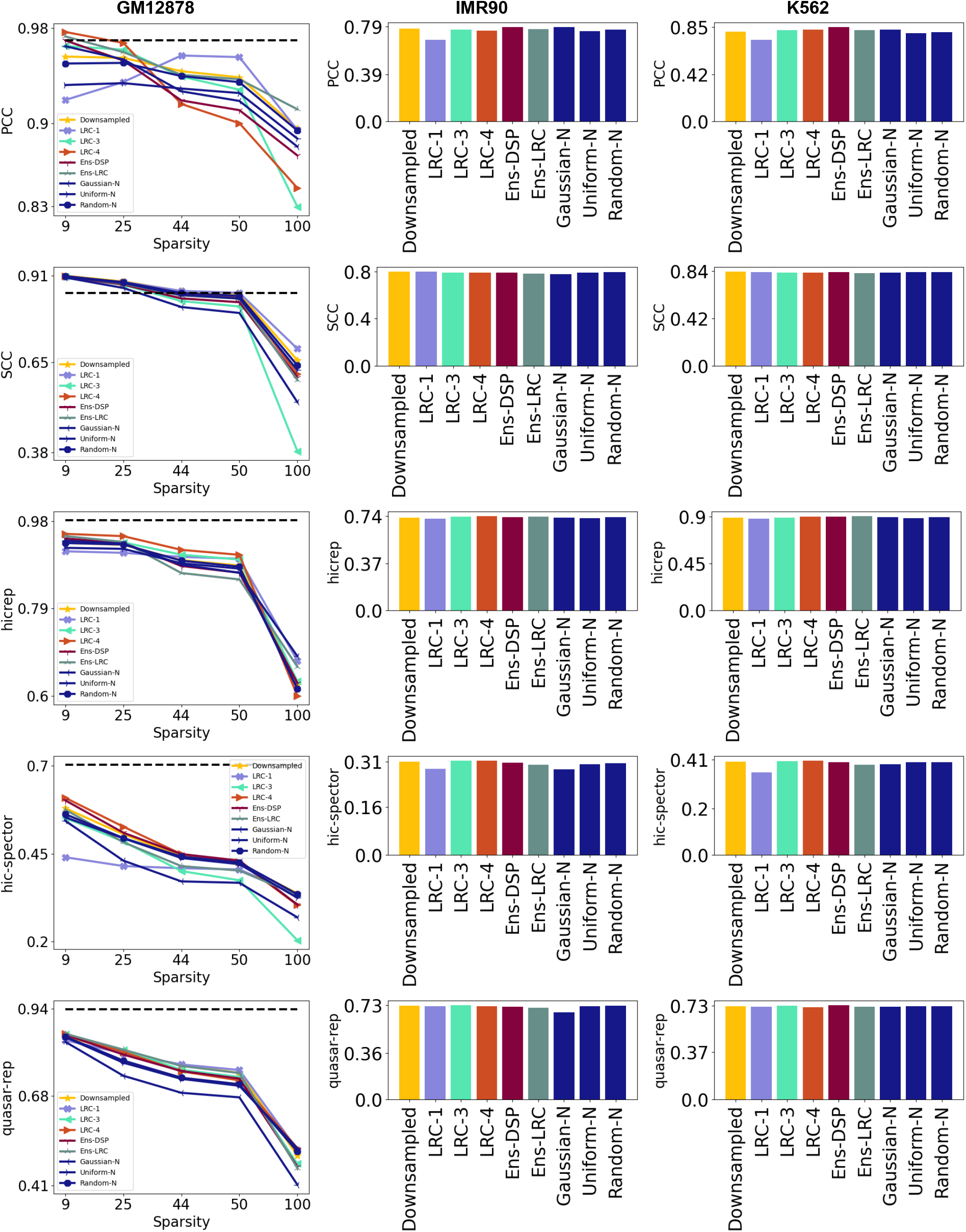
We show results of retraining across five additional metrics and also across additional retraining variants, such as downsampled datasets augmented with noise

## Footnotes

1 Synonymous with increasing the Hi-C resolution

2 synonymous with high-resolution Hi-C contact maps

3 Synonymous with low-resolution Hi-C contact map.

4 We consider an overlap if the position of both features are within *n* bin of each other. We tune value of *n* for all feature individually as summarized in the Supplementary Figure S2

